# The Flip-flop neuron – A memory efficient alternative for solving challenging sequence processing and decision making problems

**DOI:** 10.1101/2021.11.16.468605

**Authors:** Sweta Kumari, C Vigneswaran, V. Srinivasa Chakravarthy

**Affiliations:** Computational Neuroscience Lab, Department of Biotechnology, IIT Madras, Chennai, 600036, Tamil Nadu, India

**Keywords:** RNNs, LSTM, Flip-flop, Signal generation, Handwriting generation, Text generation, Video frame prediction, Action recognition

## Abstract

Sequential decision making tasks that require information integration over extended durations of time are challenging for several reasons including the problem of vanishing gradients, long training times and significant memory requirements. To this end we propose a neuron model fashioned after the JK flip-flops in digital systems. A flip-flop is a sequential device that can store state information of the previous history. We incorporate the JK flip-flop neuron into several deep network architectures and apply the networks to difficult sequence processing problems. The proposed architectures include flip-flop neural networks (FFNNs), bidirectional flip-flop neural networks (BiFFNNs), convolutional flip-flop neural networks (ConvFFNNs), and bidirectional convolutional flip-flop neural networks (BiConvFFNNs). Learning rules of proposed architectures have also been derived. We have considered the most popular benchmark sequential tasks like signal generation, sentiment analysis, handwriting generation, text generation, video frame prediction, lung volume prediction, and action recognition to evaluate the proposed networks. Finally, we compare the results of our networks with the results from analogous networks with Long Short-Term Memory (LSTM) neurons on the same sequential tasks. Our results show that the JK flip-flop networks outperform the LSTM networks significantly or marginally on all the tasks, with only half of the trainable parameters.

## Introduction

Many instances of human decision making involve integration of information over extended durations of time. Sequential processing of the information is needed in a large variety of problems, for example playing chess^1^, solving the Rubick’s cube^2^, video processing^3^, video analytics^4^, and language processing^5^. Some tasks require short-term memory of the past states. But problems arise in tasks that require long-term memory of stimulus-response relationships without loss of information.

Recurrent neural networks (RNNs), with the added feature of Long Short-Term Memory (LSTM) units, have proved capable of solving difficult sequential decision making problems^6^,^7^. RNN and LSTM combination has the ability to forget irrelevant information and retain highly discriminative information that can be held for long durations before it can contribute to the final decision. The gating mechanism of LSTM is built predominantly to overcome the issues of catastrophic forgetting and vanishing gradients that are associated with classical RNNs^8^.

Basically, there are four gates in the RNN-LSTM unit. These gates control the remembering and forgetting functions of the LSTM^6^. The input gate of the RNN-LSTM decides what new patterns of the data are to be preserved by multiplying the long-term memory with input vectors, and what patterns of the data to be forgotten by multiplying the long-term memory with forget vectors. Thus, the input gate filters out the combination of the current input data and past information stored in long-term memory and forwards it downstream. The output gate is implemented to produce the short-term memory that is used by the next LSTM unit.

We refer to deep networks that comprise LSTMs as RNN-LSTMs. A number of complex sequential tasks have been solved by using RNN-LSTMs^9^,^10^. The combination of Convolutional Neural Networks (CNNs) and LSTMs has been applied for sentiment analysis of^11^. RNN-LSTMs have been used along with a Mixture Density Network (MDN) to generate artificial handwriting samples^12^. RNN-LSTMs have produced splendid results in generating context-based text^13^,^14^,^15^,^16^. RNN-LSTM combined with the Inception architecture^17^,^18^ were used for next frame prediction in video. Attention-based RNN-LSTM networks^19^ and convolutional RNN-LSTM networks (ConvLSTM)^20^ have also been applied for human action recognition. But the main drawback of LSTM^6^ is that a single unit has 4 gating variables to be trained, resulting in high computational expense for both training and testing.

Drawing inspiration from flip-flops of digital systems, which have the ability to store memory, Holla and Chakravarthy (2016)^21^ have proposed the flip-flop neuron. It was shown how simple neuron models with bistable dynamics and/or oscillatory dynamics can be made to emulate two kinds of flip-flops – the S-R flip-flop and the Toggle flip-flop. However, since the JK flip-flop is considered to be the universal flip-flop, a neuron model fashioned after JK flip-flop is expected to deliver better results. To this end, we propose the JK flip-flop neuron which is incorporated in a deep network structure and applied to a variety of sequential decision making and sequence processing problems.

In the current study, JK flip-flop neurons are incorporated in both Convolutional (Conv) and Fully Connected (FC) layers. Learning rules of such neurons in the network were also developed. We evaluated our JK flip-flop neural networks on popular sequential tasks such as signal generation^22^, sentiment analysis^23^, handwriting generation^24^, text generation using character-based learning^14^, next frame prediction in a video^17^, lung volume prediction^25^, and action recognition^26^. RNN-LSTMs have also been applied to such tasks before, thereby making a comparative analysis possible. Furthermore, the flip-flop neurons are analogous to the bi-stable neurons in the Prefrontal Cortex (PFC), responsible for working memory functions^21^. Therefore, flip-flop neurons can be contextualized in the field of computational neuroscience as well. The proposed network of flip-flop neurons can also be considered as a “store once and retrieve forever” network^21^. This implies that once a flip-flop sets for a stimulus, the rest of the network always has access to the information that the stimulus has occurred unless it is modified by another stimulus.

## Background

RNN is one of the traditionally used networks to solve sequence processing problems. However, the inability to remember relevant information and ignore irrelevant information over long durations is one of the main drawbacks of these networks. In 1997, Hochreiter & Schmidhuber proposed the LSTM unit as a variant of vanilla RNN by adding an extra state and four gating variables. RNN-LSTMs are one of the commonly used and state-of-the-art networks to solve a wide variety of sequential problems^6,27^ like sequence-to-sequence generation, attention modelling and classifying sequences with long-term dependencies^9,11–20^,. There is also a lot of interest in studying the learning dynamics of RNN-LSTM and its memory properties^28^. Hopfield proposed^29^ a simple network with stable attractors to model memory and also applied it to solve optimization problems. Subsequently, it was also generalized to store and retrieve sequences but has serious capacity limitations for long sequences^30^. Holla and Chakravarthy^21^ used SR and Toggle flip-flops dynamics to model memory and used the network to make decisions involving long delays to demonstrate long-term memory retention. There are many networks proposed as alternatives to RNN-LSTM with a lesser number of trainable parameters, better performance and better optimization^31–34^. Although the proposed models tried to address a particular solution as an improvement to RNN-LSTM, none of them have been consistently demonstrated to be a more powerful alternative to RNN-LSTM^35^. Many deep learning libraries have used optimal unlooping of RNN-LSTM by parallelisation. However, reducing the performance bottleneck in hardware when using RNN-LSTM is still an open research question^36^.

Therefore, in the current study, we propose a JK flip-flop neuron that has dynamics and ability to preserve long-term memory. Subsequently, we exploit these dynamics in deep networks that can solve benchmark sequential problems. The results of such dynamical networks are compared with the results of the RNN-LSTM networks on the same tasks.

## Methods

**“This statement is to confirm that all methods were carried out in accordance with relevant guidelines and regulations “**

A flip-flop is an electronic component used in digital circuits to store state information. There are 4 types of flip-flops: Toggle flip- flops, D flip-flops, SR flip-flops, and JK flip-flops^27^. Using a bistable latch, a flip-flop can hold the state value of previous time steps. Along with the memory latch, based on its two inputs – Set (S) & Reset (R) – it can erase old information or latch new information. This latching property of flip-flops is utilized in digital electronics to serve as a memory unit. A single standalone flip-flop circuit can hold one bit of information. In digital systems theory, flip-flops are used to construct sequential circuits^28^. Similarly, it has been shown earlier that neurons can be designed to emulate flip-flops – the flip-flop neurons – and can be incorporated into neural architectures to solve temporal decision making problems with long delays (Holla and Chakravarthy, 2016)^21^.

### Standalone Flip-flop neuron

Although digital flip-flops are binary in nature, the flip-flop neurons in the current implementation have continuous outputs in the range of [0, 1]. The input/output function of the flip-flop helps to ensure gating property and differentiability necessary to calculate gradients for parameter updates. The JK flip-flop is the most versatile of the basic flip-flops^37^. In this study, the flip-flops are fashioned after JK flip-flops in particular since these flip-flops subsume both SR and Toggle flip-flops, and are known to be universal flip-flops^38^. Therefore, throughout the paper, the word flip-flop refers to the JK flip-flop.

The flip-flop neuron takes two competing inputs *J* and *K*, in which *J* is used to add new salient information, and K is used to remove irrelevant old information (Figure 1). Along with the two inputs, the previous state *Q*_*old*_ is also fed to estimate the state of the current time-step *Q*_*new*_. Characteristic equation of the JK flip-flop is given as:

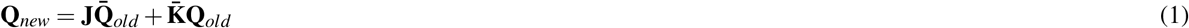

**Figure 1.**
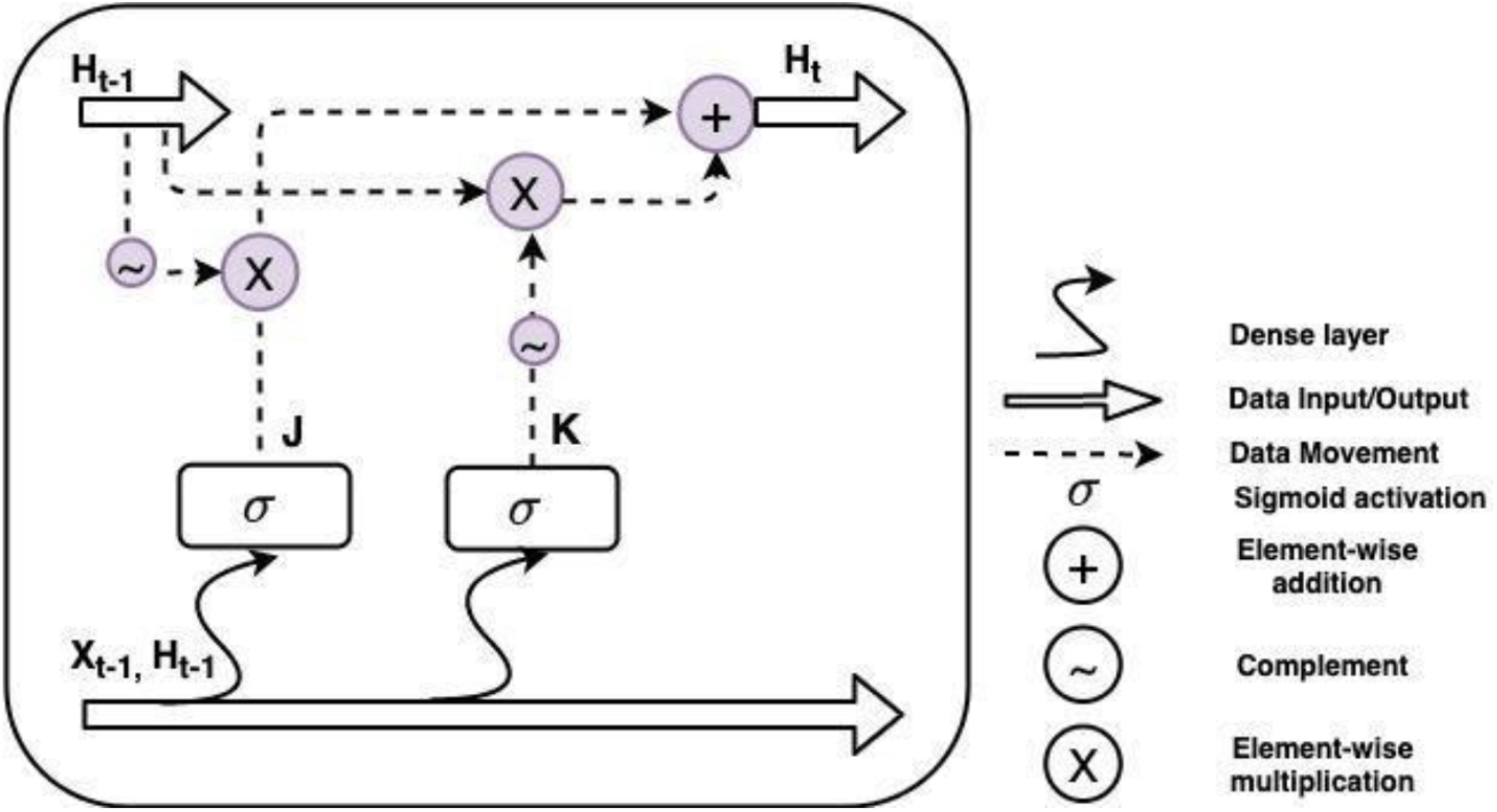
Flip-flop neuron

From Table 1, which shows the truth table of a JK flip-flop, we can see that the JK flip-flop combines the SR flip-flop and Toggle flip-flop in a single circuit. The forward propagation of the data in the flip-flop neuron is defined as,

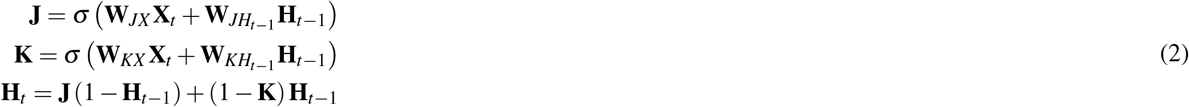

where, *σ* is sigmoid activation function, *W*′*s* are the connecting weights, and *H*_*t*_ *X*_*t*_ are the state and input values at time step ‘*t*’ respectively. Sigmoid activation and the flip-flop characteristic equation ensure all *J, K* and *H*_*t*+1_ are bounded within [0, 1].

**Table 1.** Truth table for a JK flip-flop

| J | K | $Q_{old}$ | $Q_{new}$ |
| --- | --- | --- | --- |
| 0 | 0 | $Q_{old}$ | $Q_{old}$ (no change) |
| 1 | 0 | 1 | 1 |
| 0 | 1 | 1 | 0 |
| 1 | 1 | $Q_{old}$ | $\bar{Q}_{old}$ (toggles) |

#### Training Rules

When backpropagating the loss gradients, the backward propagation across the flip-flop neuron is defined as,

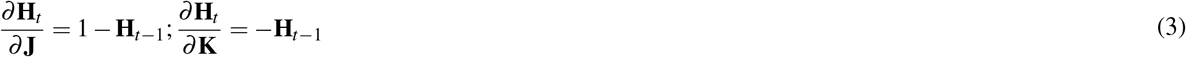

The partial derivatives with respect to **J** and **K** are back propagated to the corresponding *J* and *K* nodes.

### Flip-flop vs RNN-LSTM

Table 2 shows the comparison between flip-flop and RNN-LSTM neurons and their characteristic equations. From the forward and backward propagation equations, it is clear that the number of gating variables required for flip-flop is two (J & K) with only one recurrent state vector (H) unlike in RNN-LSTM which uses four gating variables and two recurrent state vectors. Thus, the number of trainable parameters required for flip-flops in the neural network is almost half that of RNN-LSTM. This parsimonious property of flip-flops offers many advantages including reduced training time, reduced FLOPS (floating point operations per second), faster execution in real-time and light-weight hardware realizations. Since flip-flops already have field programmable gate arrays (FPGA) and other hardware realizations, such methods can be used in design of hardware realizations for flip-flop neural networks^39^.

**Table 2.** Comparison between the architecture and characteristic equations of flip-flop and RNN-LSTM neuron

| Flip-flop neuron | LSTM neuron |
| --- | --- |
| $J = \sigma(W_{JX}X_t + W_{JH_{t-1}}H_{t-1})$ $K = \sigma(W_{KX}X_t + W_{KH_{t-1}}H_{t-1})$ $H_t = J(1 - H_{t-1}) + (1 - K)H_{t-1}$ | $I_t = \sigma(W_{IX}X_t + W_{IH_{t-1}}H_{t-1})$ $F_t = \sigma(W_{FX}X_t + W_{FH_{t-1}}H_{t-1})$ $O_t = \sigma(W_{OX}X_t + W_{OH_{t-1}}H_{t-1})$ $\tilde{C}_t = \tanh(W_{GX}X_t + W_{GH_{t-1}}H_{t-1})$ $C_t = \sigma(F_t C_{t-1} + I_t \tilde{C}_t)$ $H_t = \tanh(C_t) O_t$ |

In network realizations, flip-flop neurons can be organized in two ways: flip-flop neurons in the convolutional layer (named as “convolutional flip-flop layer” or ConvFF), and flip-flop neurons in the fully-connected layer (named as “fully connected flip-flop layer” or FCFF).

### Fully connected flip-flop layer (FCFF)

In digital logic circuits, it is known that arrays of combinatorial logic are followed by sequential units^40^, which enables the circuit to utilize the inputs and present appropriate state representation to the memory units. Analogously, we construct neural networks in which the input is first passed through a feedforward stage before it is presented to a layer of flip-flop neurons. Beyond the flip-flop layer, we need one or more hidden layers that transform the outputs of the flip-flop neural layer into the desired output.

### Convolutional flip flop layer (ConvFF)

The only difference between a FCFF layer and a convolutional flip-flop layer is that in the latter the flip-flop neurons are organized as 2D arrays with convolutional receptive fields (shown in Figure 2). We use a convolutional flip-flop neural network (ConvFFNN) for sequential tasks that involve images such as next frame prediction in video, predict lung volume in sequence of CT scan images, and action recognition.

**Figure 2.**
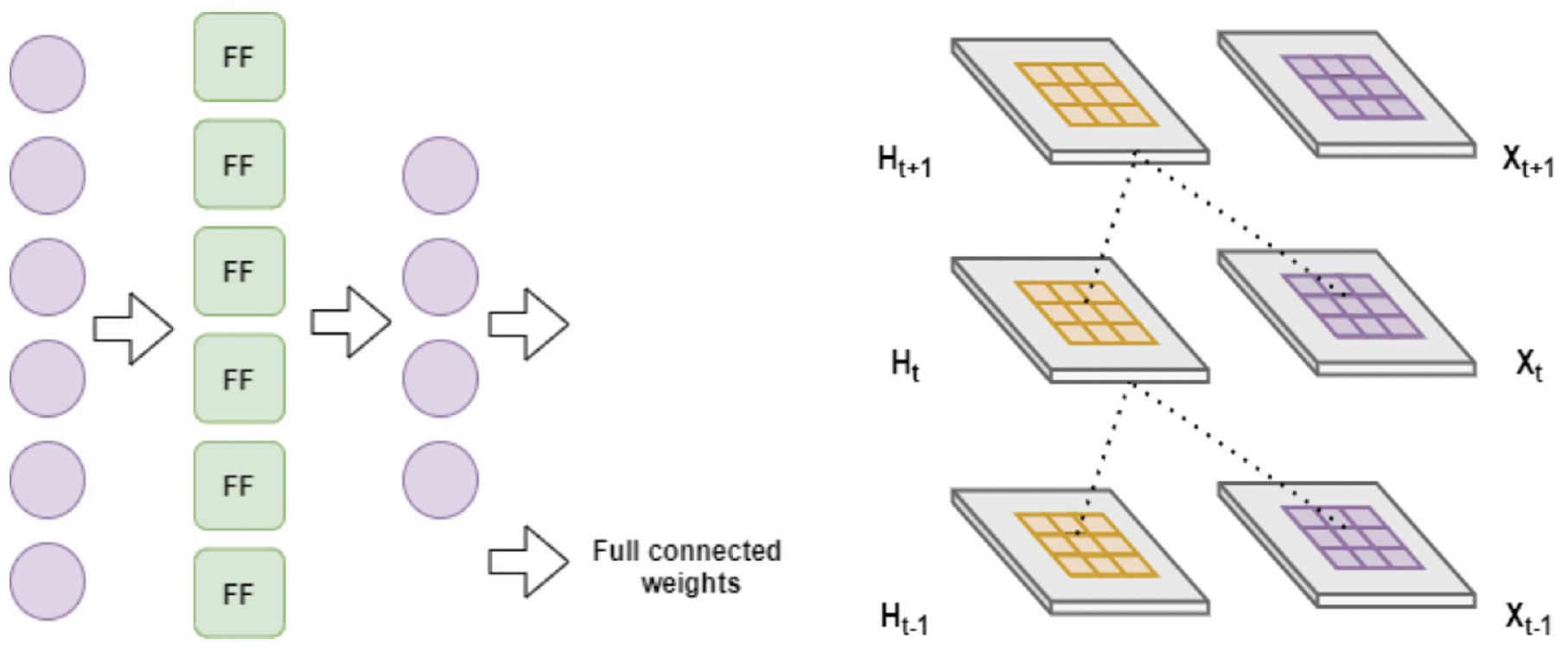
General network architecture of FCFF and ConvFF

## Experiments and Results

In this section we compare the performance of the flip-flop neural networks and RNN-LSTM in a variety of sequential tasks. Depending on the task demands we model two versions of flip-flop networks. FCFF layers are used in case of vector-based sequential tasks whereas ConvFF layers are applied for image based sequential tasks. Seven different tasks are considered to make the performance comparison reasonably comprehensive. Comparison in terms of memory requirements in the Central Processing Unit (CPU), Graphics Processing Unit (GPU) and Tensor Processing Unit (TPU) is also presented. All the implementations are carried out in Python 3 and Tensorflow2 is used for implementing and training network models. Source code for all the experimentations mentioned in this section is available in this link.

### One dimensional sequential tasks

#### Signal generation

Signal generation is a standard one-to-many sequence generation problem, where the network is trained on n-different types of labeled signals. When a specific label is given as the input, the network generates the corresponding signal.

For this task, two datasets are used. In the first dataset, two classes of signals made of sinusoidal functions with arbitrary parameter (amplitude, frequency and initial phase) values are generated synthetically with each class having 64 signals of 100 time steps each. Noise from Gaussian distribution *G*(0, 1) is added to the data to differentiate among individual signals of the same class. First row of Figure 3 shows the sample signals from two classes of dataset 1 and the input label is a one-hot-encoder string of size 3. Similarly, for the second dataset, the whole data generation process is repeated by replacing sinusoidal functions with sampling random data points from uniform distribution *U* (−1, 1). In this dataset, the data points do not show any periodic behavior and therefore, the model needs to memorize all the data points of the two different classes of sequence. The second row of Figure 3 shows the sample signals from two classes of dataset 2.

**Figure 3.**
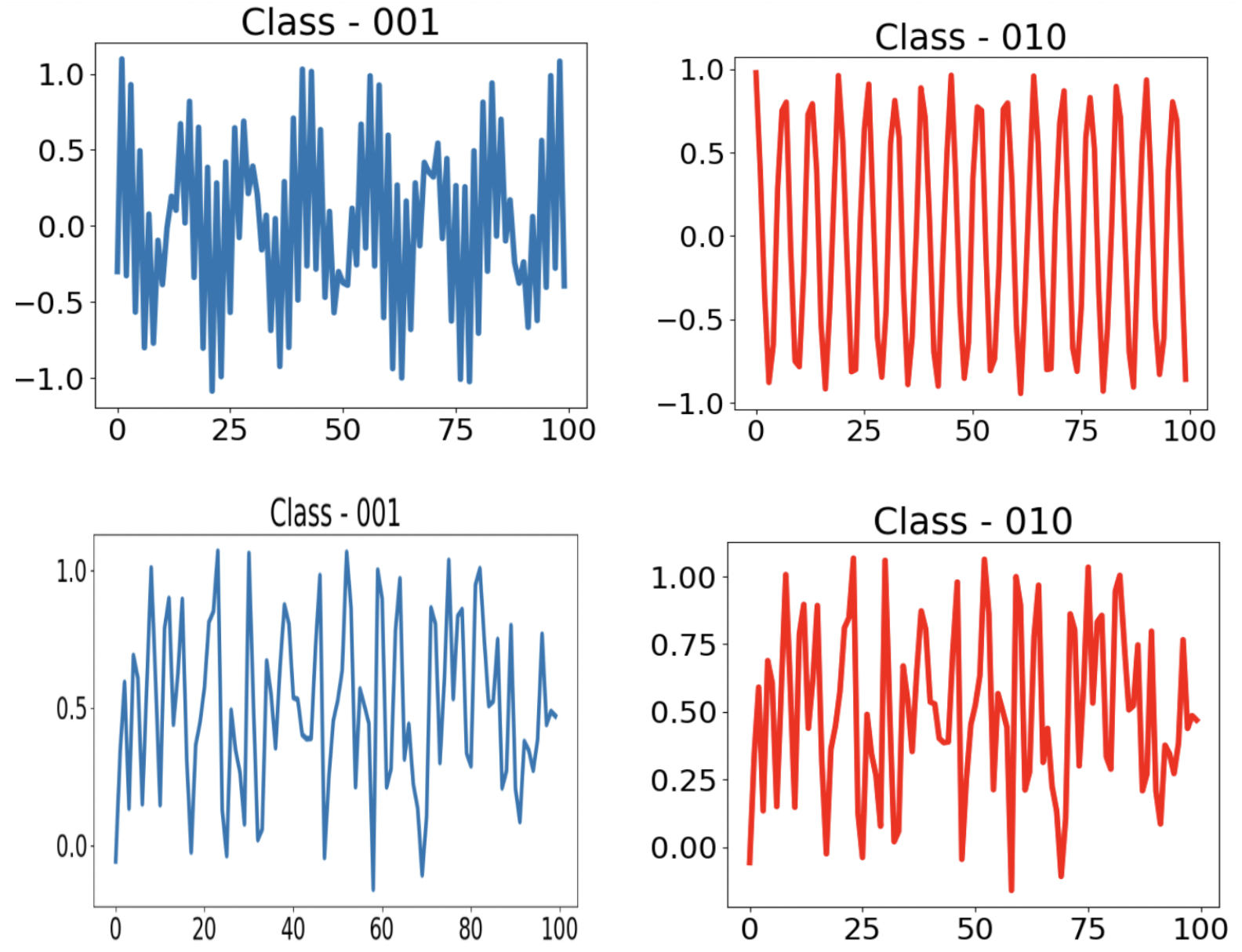
Sample of synthesized signals made of two classes from dataset 1 and 2 are shown in first and second row respectively.

The network for this task has two FCFF layers of 50 neurons, followed by a dense layer with a single neuron. Mean Squared Error (MSE) and Adam (learning rate = 0.001) are configured as the error criterion and optimizer respectively. The same architecture and setup is used for RNN-LSTM, replacing flip-flop neurons with RNN-LSTM neurons.

Table 3 shows the comparison between the above two networks. Figure 4 and 5 show the generated signals by FFNN and RNN-LSTM respectively. It is clear that signals generated by RNN-LSTM miss a few points (Figure 5) in both the classes whereas FFNN generated more accurate signals (Figure 4). The same observation is also reflected in their MSE loss (Table 3).

**Table 3.** Comparison of flip-flop network and RNN-LSTM in Signal generation

|  | FFNN | RNN-LSTM |
| --- | --- | --- |
| MSE Loss - Dataset 1 | $1.341 \times 10^{-5}$ | $3.731 \times 10^{-4}$ |
| MSE Loss - Dataset 2 | $2.394 \times 10^{-4}$ | 0.0229 |
| Number of trainable parameters | 15,751 | 31,051 |
| Number of Epochs | 2000 | 2000 |

**Figure 4.**
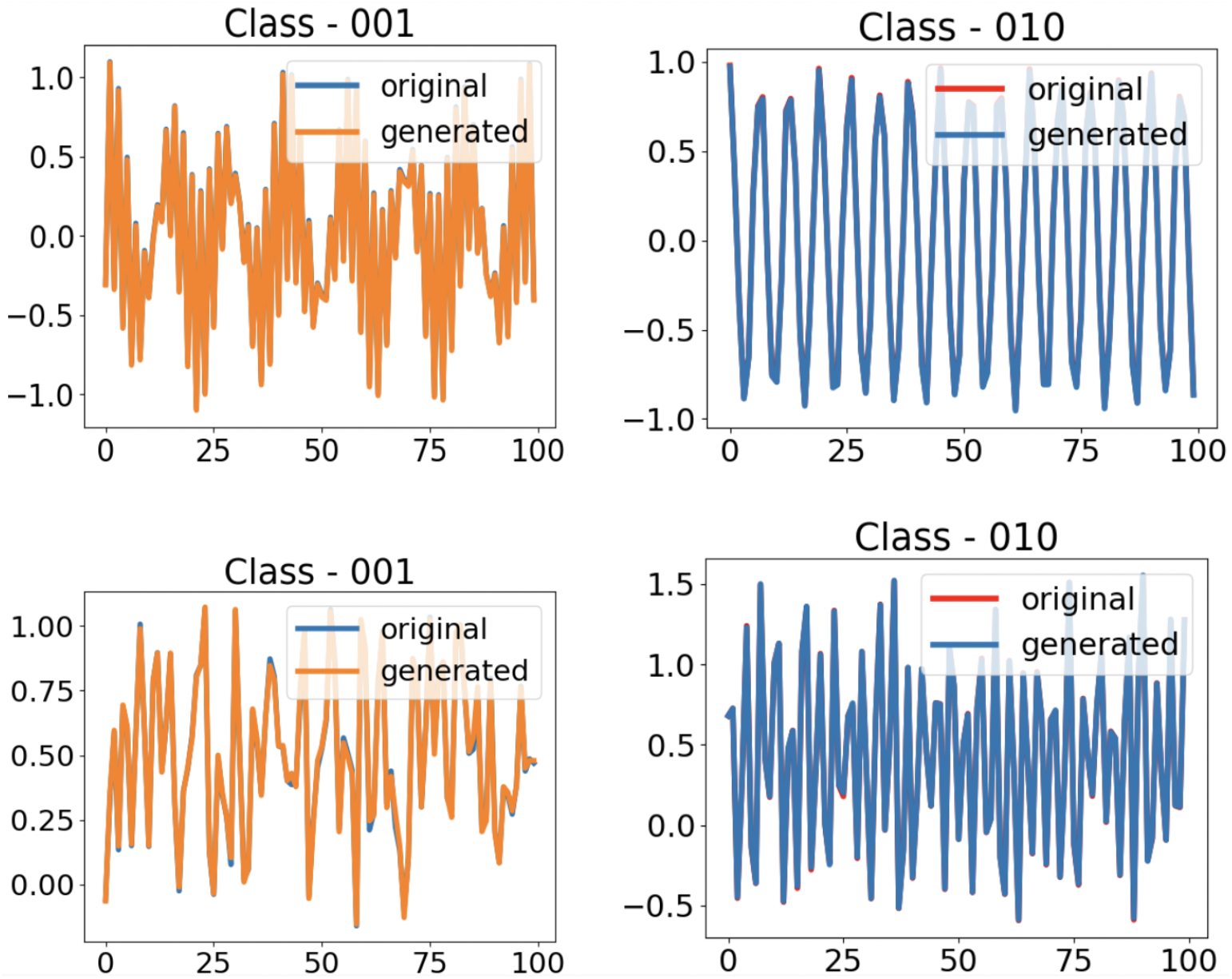
Signals generated (Dataset 1 and 2 in first and second row of the figure respectively) by the FFNN. Since the original and generated signals perfectly overlap they are not easily distinguishable in the plot

**Figure 5.**
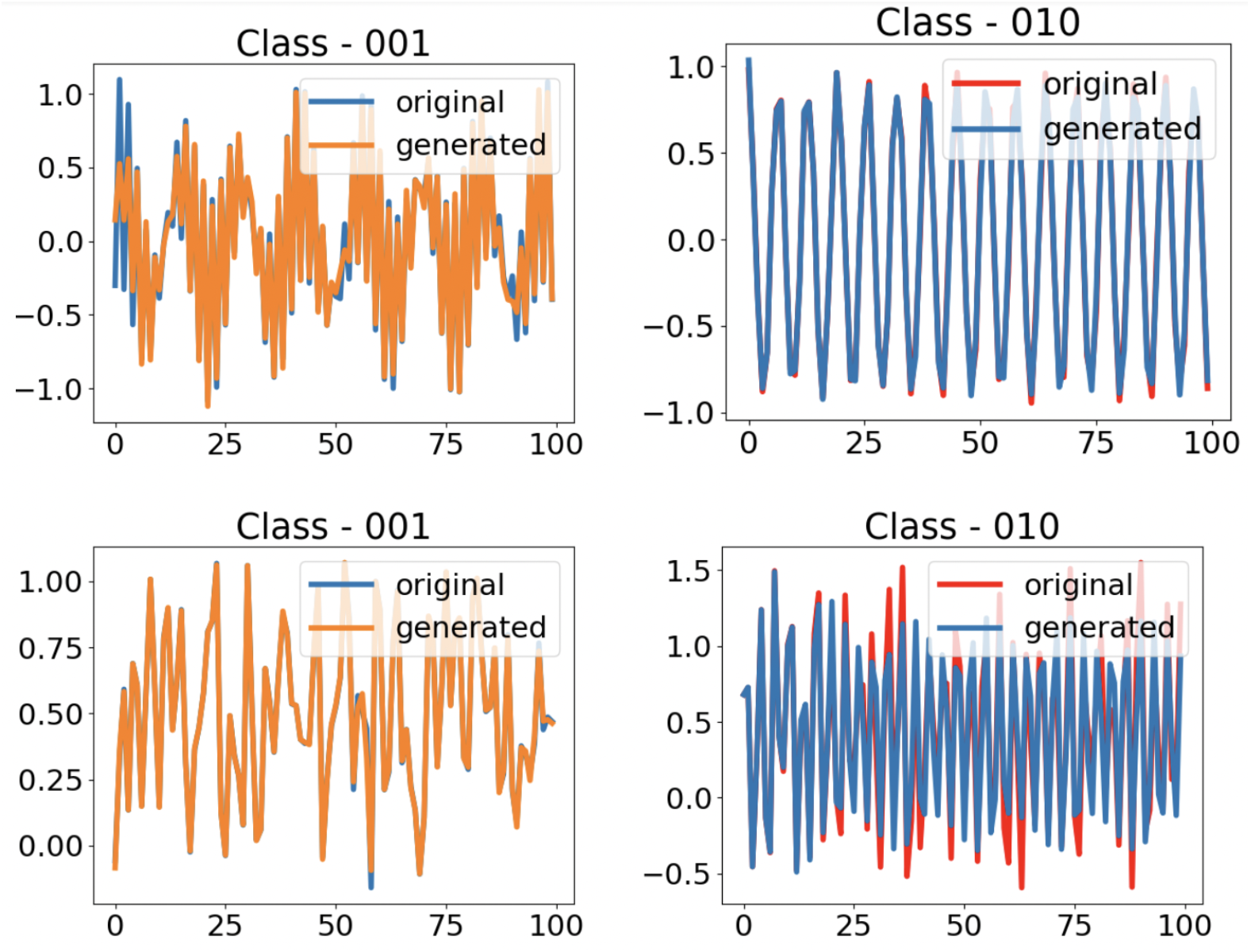
Signals generated by RNN-LSTM (Dataset 1 and 2 in first and second row of the figure respectively)

#### Sentiment analysis

This task was chosen to showcase the ability of the FFNN to classify sequences. Over the period of training, the model should learn the discriminatory features from the data to classify whether the sequence of words represents a positive or a negative review. Movie review sentiment analysis is a two-class (positive/negative) classification problem. IMDB’s large movie review dataset **??** is often used for sentiment analysis, where maximum review length is set as 500 words. The dataset provides separate datasets for training and testing, with 25, 000 polar movie reviews in each set. While training, the training set is further split into training and validation in a 7 : 3 ratio.

The proposed network architecture is shown in Figure 6. The words are encoded and passed to the embedding layer followed by a bidirectional FCFF (BiFCFF) layer with 100 units and a sigmoidal neuron in the last layer. Binary Cross Entropy (BCE) is used as the loss function with Adam (learning rate = 0.001) as the optimizer. Flip-flop cells are replaced with RNN-LSTMs in the recurrent part to obtain the RNN-LSTM version of the model.

**Figure 6.**
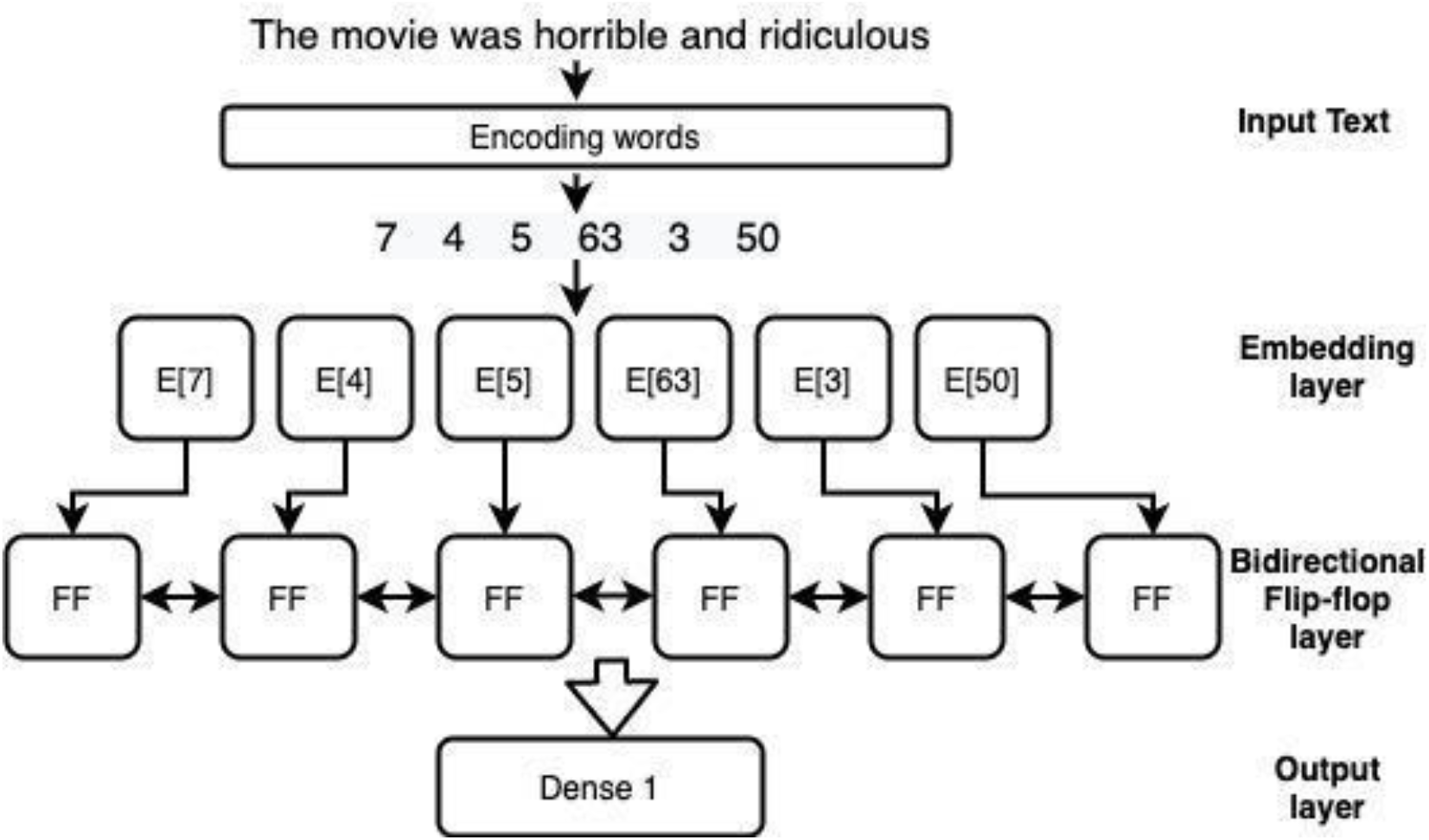
BiFFNN architecture for sentiment analysis task

Table 4, shows comparison between two networks in terms of accuracy, loss and number of trainable parameters. RNN- BiLSTM performs (with accuracy 85.19%) only slightly better than BiFFNN (with accuracy 85.07%).

**Table 4.** Comparison of performance between BiFFNN and RNN-BiLSTM for sentiment analysis task

|  | BiFFNN | RNN-BiLSTM |
| --- | --- | --- |
| Validation BCE | 0.3596 | 0.3433 |
| Training Accuracy | 86.01% | 86.24% |
| Validation Accuracy | 85.07% | 85.19% |
| Number of trainable parameters | 53,600 | 106,400 |

#### Handwriting generation

Handwriting generation is a challenging problem, and unlike prediction, the model is not constrained by any order of letters nor number of points for a letter. The trained handwriting generation neural network should exhibit smooth transitions from one letter to the next, maintain the baseline, and generalize different styles.

The dataset used for this task is, IAM online handwriting dataset^41^. The dataset is originally in XML format with 13, 049 text lines, 86, 272 word instances and 11, 059 unique words contributed by 221 writers. Each coordinate point of a character is three-dimensional, representing: *x* & *y* coordinates and end of stroke signal.

Figure 7 shows the handwriting generation network architecture. The neural network contains a stack of three flip-flop layers with each layer getting input from the layer below followed by Mixed Density Network (MDN)^42^ in the last layer. The MDN network is used to output parameters of the bivariate Gaussian model to estimate negative log-likelihood loss. The window layer is used as an attention unit that gives the weighted output of input character identifiers. The output from the last layer is a three-dimensional tuple representing *x, y* coordinates and end of stroke bit, which in turn is fed back as input for the next time step. The window layer gives three output parameters, *a, b* and *c* that represent importance, width and position of the window respectively. When a L-long input sequence of character identifiers 0 ≤ *c* ≤ *L* is given as input to generate T-long points 0 ≤ *t* ≤ *T*, the window weight *W* (*t, c*) is calculated by convolving with K- Gaussians.

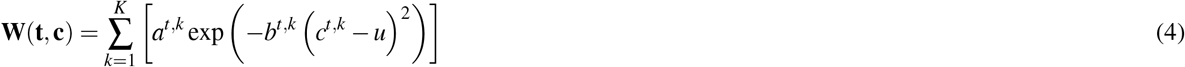

**Figure 7.**
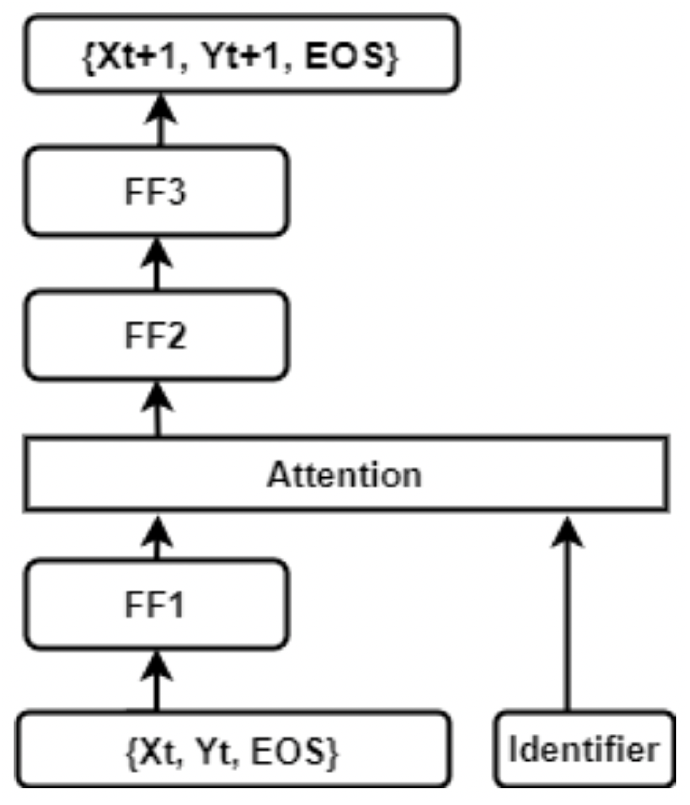
FFNN architecture of handwriting generation

After the training, sampling is done with a specific bias value to vary the level of diversity. As the bias increases, the generated handwriting looks identical for any number of trials and more legible. The network is configured with 400 flip-flop units at all three layers and sequence length of 200 is used for input data. A set of 20 mixture bivariate Gaussians, Adam optimizer (learning rate = 0.001) and negative log loss function are used for training.

Table 6 shows the handwriting generated by both the networks at different values of biases. It is clearly visible that for larger bias values, the generated handwriting samples look more legible. However, it is conclusive that the generated handwriting of flip-flops is more legible than RNN-LSTM at all levels of bias values. Also, Table 5, tabulates the training loss and epochs by both the models.

**Table 5.**
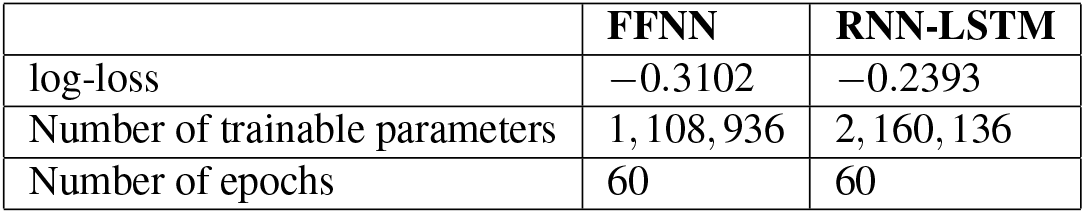
Performance comparison for handwriting generation task

**Table 6.**
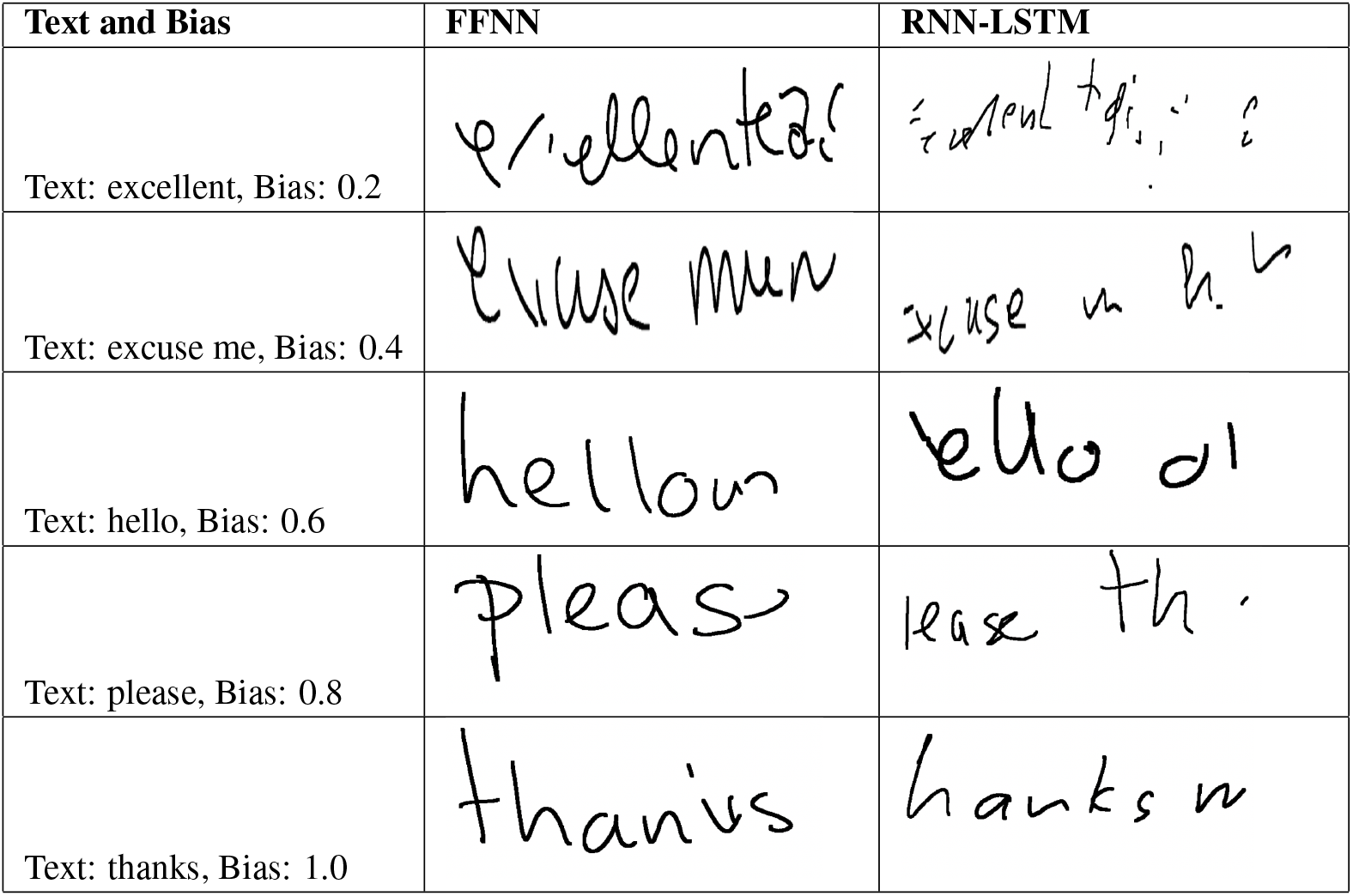
Generated handwritings of FFNN and RNN-LSTM with different biases

#### Text generation using character-based learning

The task of text generation consists of generating text that reads natural and is indistinguishable from human-generated text. Information is being created everywhere in social media in large volumes for example, Twitter tweets, WhatsApp messages, Facebook posts. A good part of the information exists in textual form, which is highly unstructured in nature.

In order to extract knowledge from this vast text corpus, methods from Natural Language Processing^43^ (NLP) and deep learning are being used extensively. Deep learning literature offers many algorithms in the field of NLP like RNNs (for sequence modeling), RNNs with external memory (RNN-EM) (to improve memory capacity of RNN)^44^, Gated Feedback RNNs (GF-RNN) (stacking multiple recurrent layers with gating units)^45^, Conditional Random Fields as RNNs (CRF-RNN) (for probabilistic graphical modeling)^46^, Quasi-RNNs (Q-RNN)(using parallel time-steps for sequence modeling)^47^, Memory Networks (for Question Answering (QA))^48^, and Augmented Neural Networks (Neural Turing Machines)^49^. Popular deep learning models known as Variational Auto-Encoders (VAEs) and Generative Adversarial Networks (GANs)^50^ which has revolutionized the field of computer vision particularly image generation (CycleGAN - for cross-domain transfer), have also been used for text generation^51^. In the current study, we used FFNN unlike the previous studies to show the long and short term memory capacity of the flip-flop neurons comparative to the RNN-LSTM. In this experiment, character sequences from Shakespeare’s poetry have been used to train the FFNN and RNN-LSTM to predict the next character in the given sequence. After training, longer sequences of text can be generated by running the model recursively.

Total length of the text in Shakespeare’s poem is 1, 115, 394 characters where 65 unique characters are present. The proposed FFNN takes input sequences where each sequence is of 100 characters. Input sequences are passed through one embedding layer of size 66 × 256, where 66 is the word size and 256 is the embedding dimension. Output from the embedding layer is passed through one FCFF layer of 1024 neurons. The output of the FCFF layer is passed through one fully connected layer containing 66 output linear neurons to predict the next character of the sequence. Batch size, epochs, and learning rate is taken as 512, 100, 0.01 respectively. Sparse cross-entropy loss^52^ is computed and back propagated using the Adam optimizer. We also trained the RNN-LSTM network with the same number of layers and neurons just by replacing the flip-flop neural units with the LSTM units. The same set of dataset and values of hyper-parameters were used for training the RNN-LSTM network. A comparison of the SCE loss of both of the recurrent networks in is shown in Figure 8, and Table 7. Number of trainable parameters and time required per epoch in case of FFNN is lesser than RNN-LSTM. Generated texts are shown in Table 8, when input word ROMEO is given to both the networks. From the table, it can be seen that the number of misspelled words (are highlighted with bold fonts in the Table 8) is higher in case of RNN-LSTM than FFNN.

**Table 7.**
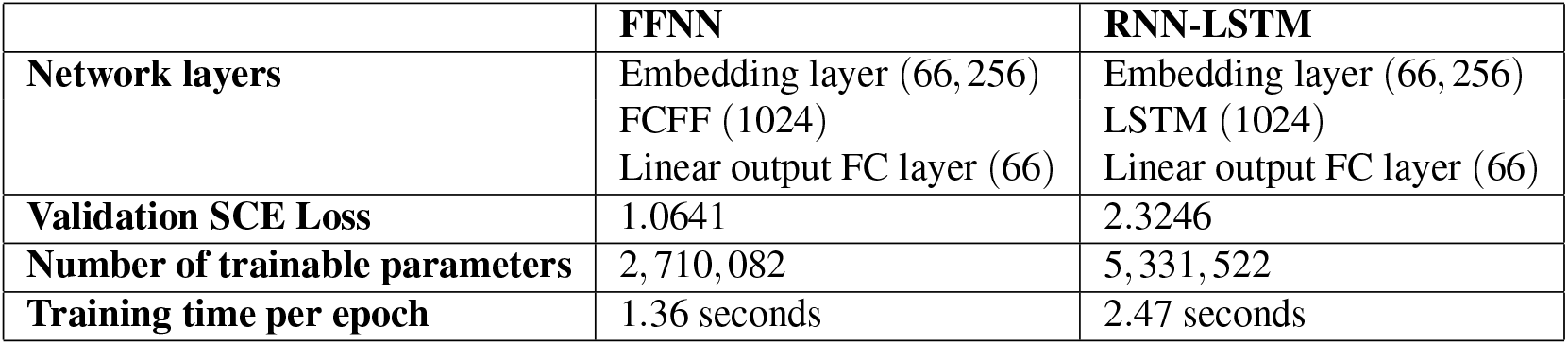
Network configuration of both FFNN and RNN-LSTM, results in terms of sparse cross entropy (SCE) loss, number of trainable parameters and training time required for each epoch are shown in the table

**Table 8.**
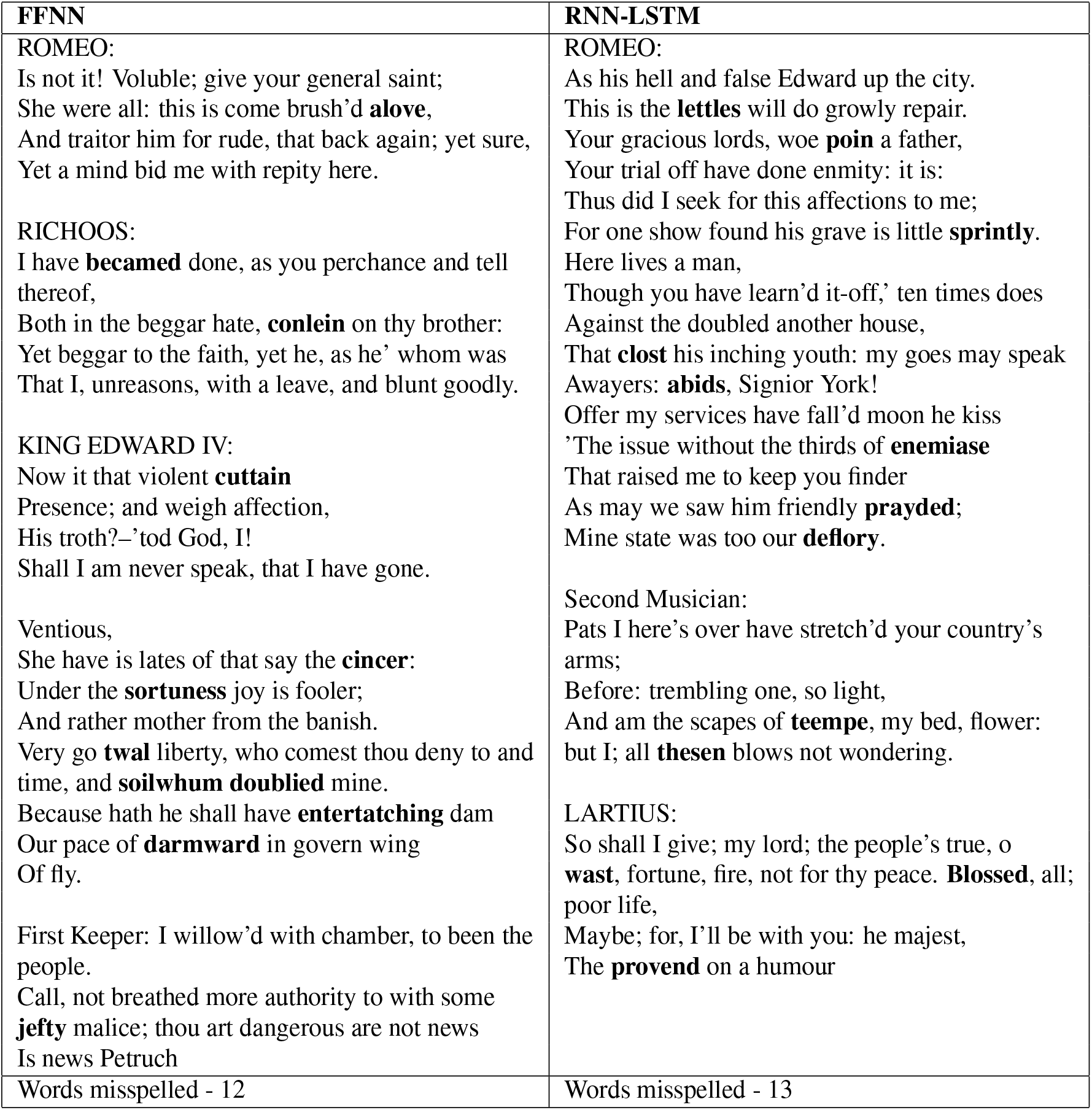
Text generated (1000 characters) when input ROMEO is given. Misspelled words are highlighted

**Figure 8.**
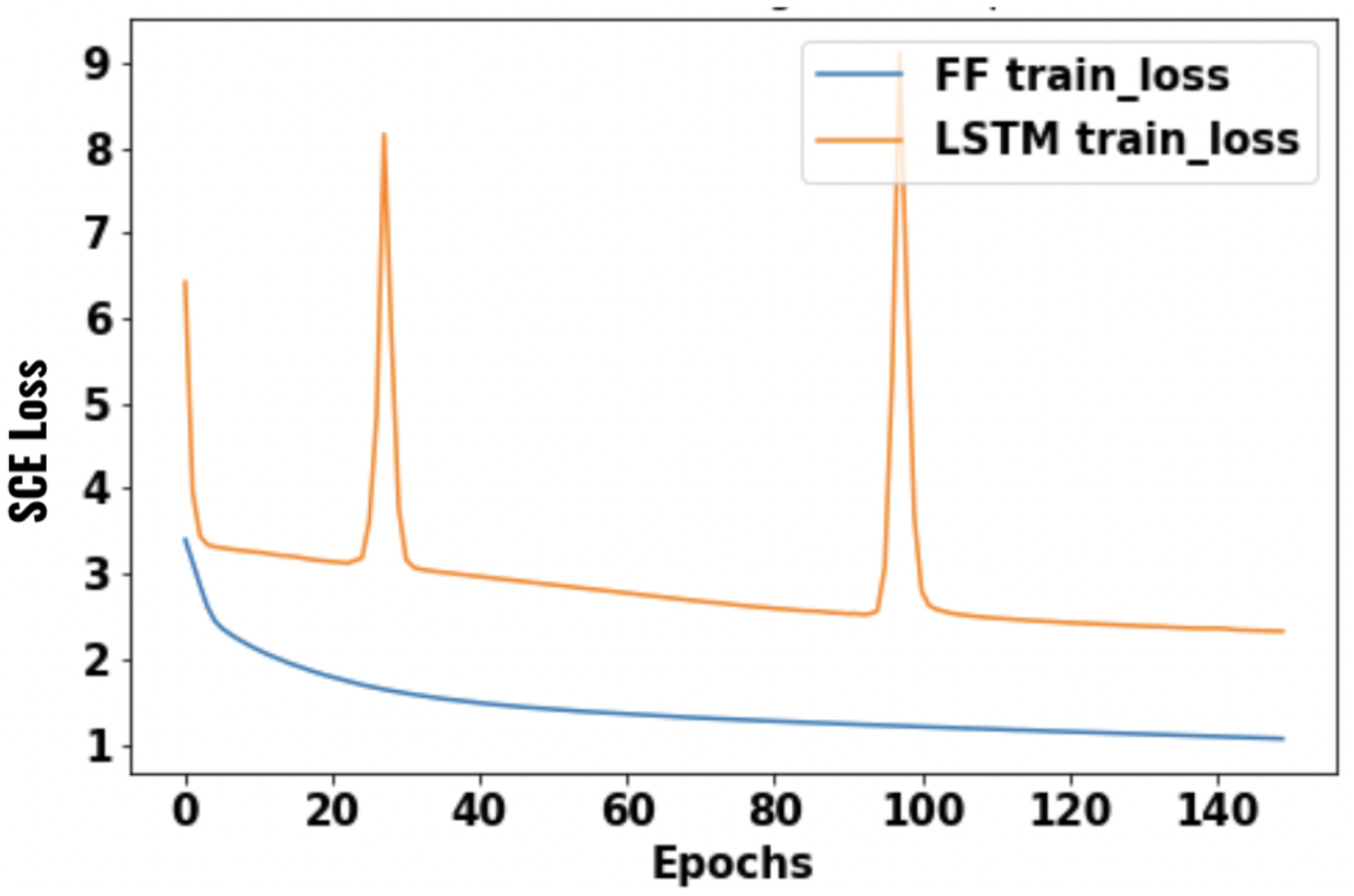
FFNN vs RNN-LSTM - Training loss vs Epochs

#### Image based sequential task

In the video frame prediction task, the network has to predict the frame at time ‘*t* + 1’, given the frame at time ‘*t*’. A synthetic dataset was used to conduct this experiment. The data consists of short video clippings where each video has 16 black and white frames of size 40 × 40. The frame consists of moving squares in one of all possible directions in 360 degrees around the square. All squares move in the same direction. The size of the squares is in the range of 2 to 4 pixels (Figure 11). Each video in the data contains 15 frames. 1000 videos were generated in total, out of which 800 videos were used for training and 200 videos for validation. 16 videos in one batch wherein the first 15 frames (frames from *t* = 0 to *t* = 14) in each video were fed into the network, and the network has to predict the next 15 frames (frames from *t* = 1 to *t* = 15).

#### Video Frame Prediction

The details of the proposed convolutional flip-flop neural network (ConvFFNN) architecture are shown in Table 9. The network is trained by backpropagating the binary cross-entropy loss between the 2-dimensional output of the network and the next frame ground truth using Ada-delta optimizer. Training is done for 100 epochs.

**Table 9.** Network configuration of both ConvFFNN and RNN-ConvLSTM for next frame prediction and results in terms of accuracy, loss, and number of trainable parameters are shown here

|  | ConvFFNN | RNN-ConvLSTM |
| --- | --- | --- |
| **Network layers** | 4× ConvFF2D, 3 × 3,40 filters,<br>Batch-norm<br>1× Conv3D, 3 × 3 × 3, 1 filter | 4× ConvLSTM2D, 3 × 3,40 filters,<br>Batch-norm, dropout-0.3<br>1× Conv3D, 3 × 3 × 3, 1 filter |
| **Validation accuracy** | 99.75% | 95.42% |
| **Validation BCE Loss** | 0.0543 | 0.2536 |
| **Number of trainable parameters** | 204,041 | 406,681 |
| **Training time per epoch** | 1.36 seconds | 2.47 seconds |

To compare the performance of the proposed ConvFFNN, we created a RNN of convolutional LSTM units (RNN- ConvLSTM). The RNN-ConvLSTM neural network is identical to the ConvFFNN network; the only difference is that 4 layers of ConvFF2D are replaced with ConvLSTM2D layers (shown in Table 9). Dropout with a keep probability of 0.3 is also applied after each ConvLSTM2D layer whereas it is not applied in the ConvFFNN network. The RNN-ConvLSTM network is also trained using the same loss function and the same optimizer as the ConvFFNN network. But in the RNN-ConvLSTM network, the total number of trainable parameters is 406,681, which is almost twice the number of the trainable parameters in the ConvFFNN network. This makes the ConvFFNN network almost 2 times faster than the RNN-ConvLSTM network (shown in Table 9. After training both networks, RNN-ConvLSTM gives the prediction with validation loss of 0.2536 whereas ConvFFNN gives the prediction with validation loss of 0.0543, shown in Table 9 (Figure 10). The validation accuracy given by RNN-ConvLSTM is 95.42% whereas it is 99.71% by ConvFFNN (Table 9 and Figure 9). Predicted frames are shown in Figure

**Figure 9.**
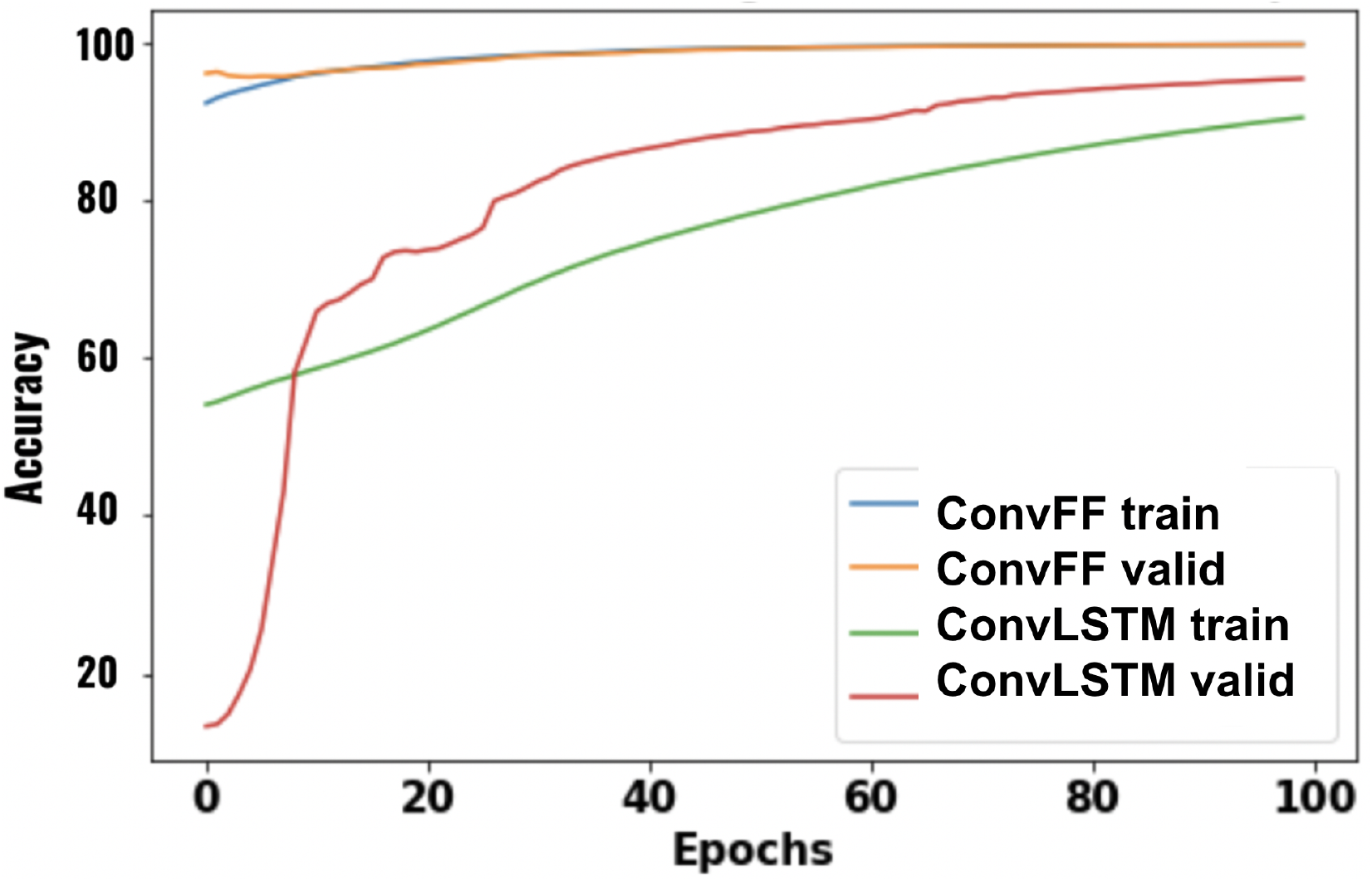
ConvFFNN vs RNN-ConvLSTM - Training and Validation Accuracy vs Epochs

**Figure 10.**
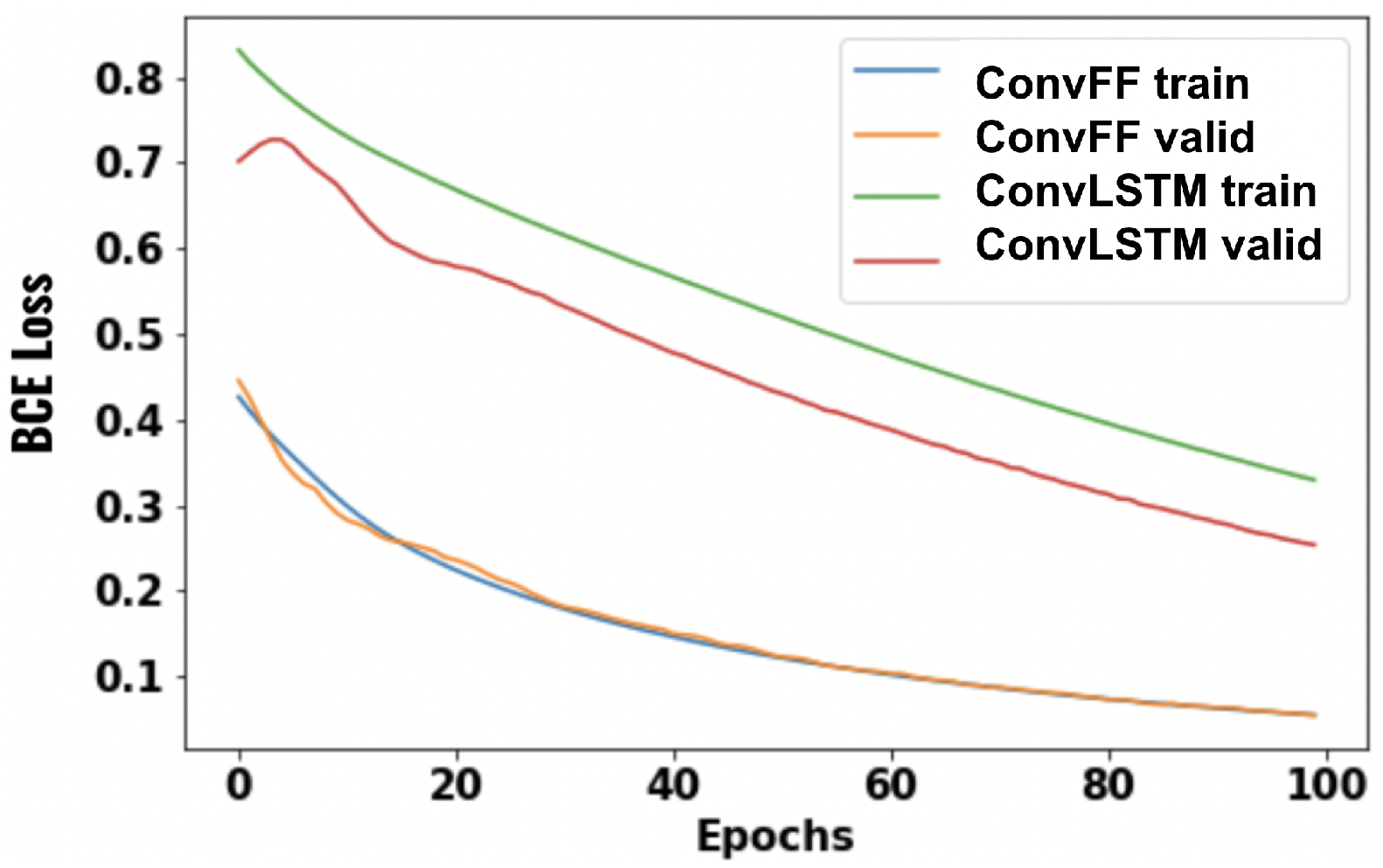
ConvFFNN vs RNN-ConvLSTM - Training and Validation Loss vs Epochs

**Figure 11.**
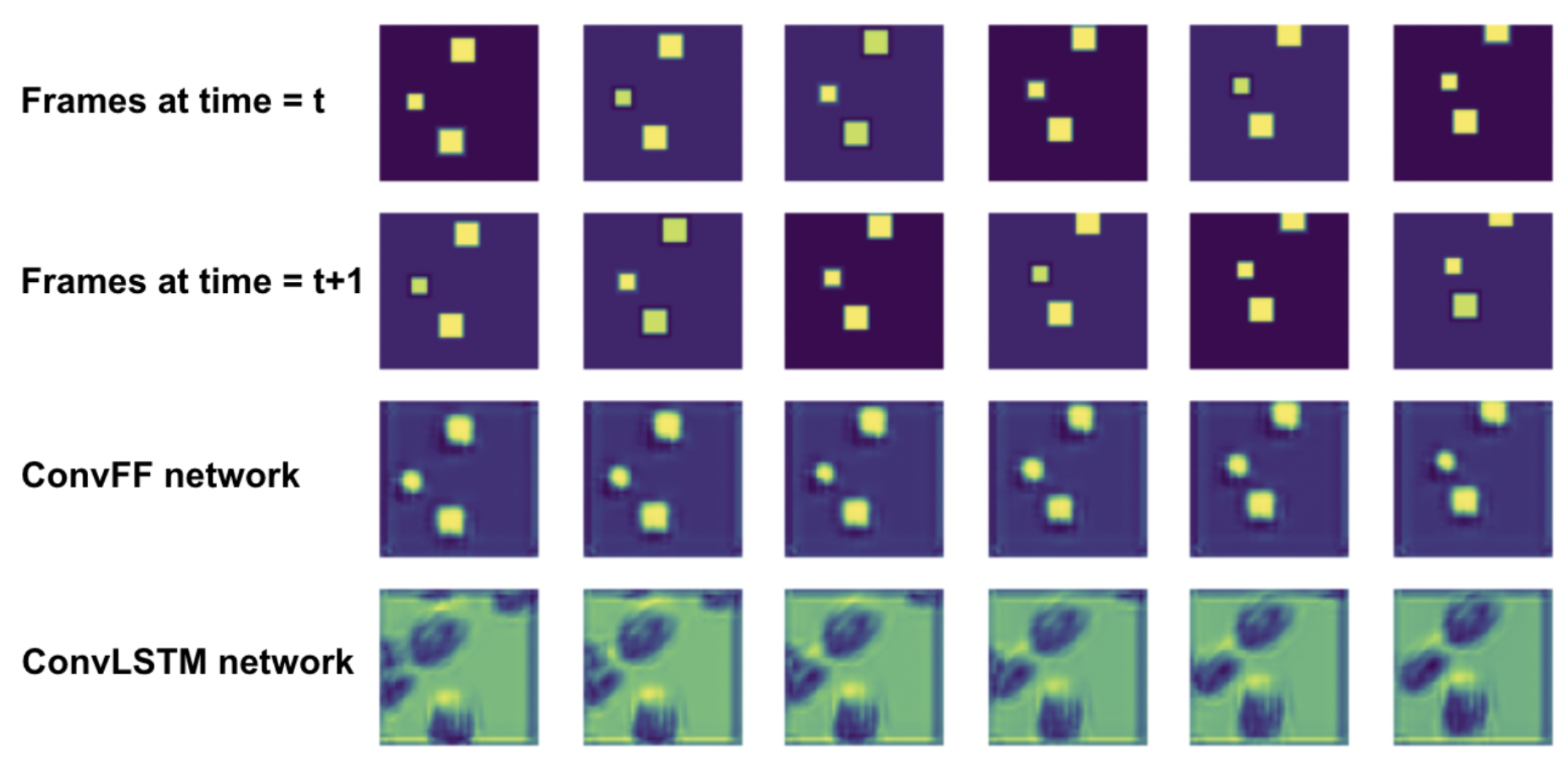
ConvFFNN vs RNN-ConvLSTM - Prediction results are shown in last two rows 11, where it is clearly demonstrated that ConvFFNN is able to predict the frames much better than the frames predicted by RNN-ConvLSTM netowrk.

#### Lung volume prediction in CT scans

The Computed Tomography (CT) scan of the lung consists of a sequence of frames, corresponding to sections of the lung parallel to the horizontal plane. The problem of estimating the lung volume from the CT scan consists of integrating the volumetric information spread out over the sequence of frames. To accomplish this task, the model is trained on a sequence of CT scans images from 100 lungs to predict the lung volume. Before training, the images are augmented with parameters like rotation range 15, width shift range 0.15, height shift range 0.15, shear range 0.1, zoom range 0.25, fill mode ‘nearest’, horizontal flip True, and vertical flip False. Width and height of the scanned images is 64. Totally 790 images in the training set and 418 images in the validation set were used.

The detailed architecture of the proposed bidirectional convolutional flip-flop neural network (BiConvFFNN) is given in Table 10. ReLU activation function is used in each of the hidden layers and sigmoid activation function is used in the output layer. To compare the results with the RNN-LSTM neuron, we implemented RNN of bidirectional convolutional LSTM (RNN-BiConvLSTM) with exactly the same layers by replacing the BiConvFF2D layers with bidirectional ConvLSTM2D (BiConvLSTM2D) layers. Both the networks are trained by backpropagating the binary cross-entropy loss^53^ using the Adam optimizer. Training is done for 20 epochs with batch size 16. Achieved loss of 0.0797 in BiConvFFNN is insignificantly less than the loss of 0.0936 in RNN-BiConvLSTM, shown in Table 10. The same table also shows that the number of training parameters of BiConvFFNN is almost half than the number of training parameters of RNN-BiConvLSTM. Figure 12 depicts the plot of training and validation loss with epochs for both the networks. Predicted lung volume of two samples (one row for one sample) by both the networks (third and forth columns) is shown in Figure 13 with their corresponding ground truth lung volume (second column).

**Table 10.**
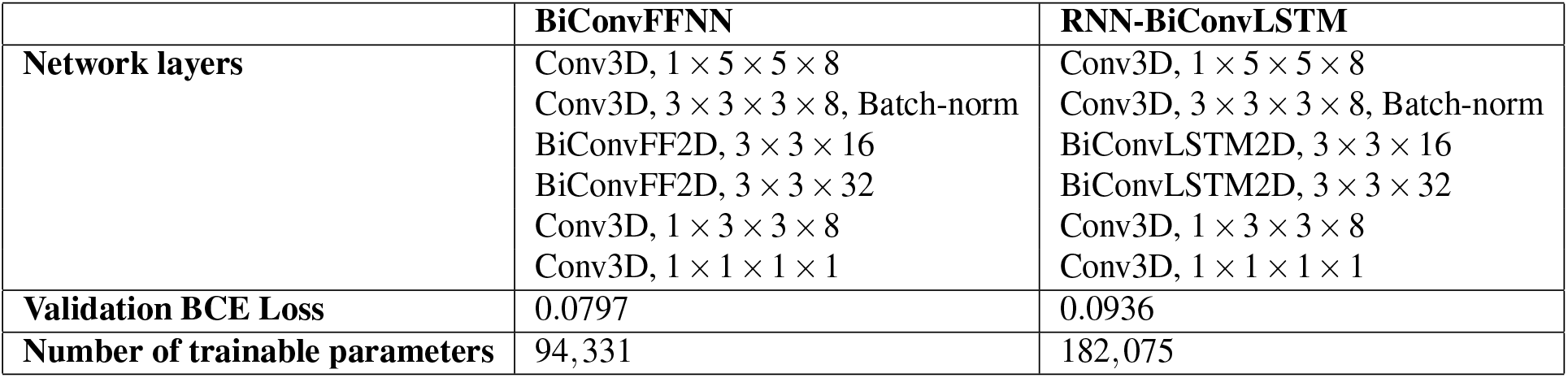
Network configuration of both BiConvFFNN and RNN-BiConvLSTM for lung volume prediction and results in terms of loss, and number of trainable parameters are shown here

|  | **BiConvFFNN** | **RNN-BiConvLSTM** |
| --- | --- | --- |
| **Network layers** | Conv3D, $1 \times 5 \times 5 \times 8$<br>Conv3D, $3 \times 3 \times 3 \times 8$, Batch-norm<br>BiConvFF2D, $3 \times 3 \times 16$<br>BiConvFF2D, $3 \times 3 \times 32$<br>Conv3D, $1 \times 3 \times 3 \times 8$<br>Conv3D, $1 \times 1 \times 1 \times 1$ | Conv3D, $1 \times 5 \times 5 \times 8$<br>Conv3D, $3 \times 3 \times 3 \times 8$, Batch-norm<br>BiConvLSTM2D, $3 \times 3 \times 16$<br>BiConvLSTM2D, $3 \times 3 \times 32$<br>Conv3D, $1 \times 3 \times 3 \times 8$<br>Conv3D, $1 \times 1 \times 1 \times 1$ |
| **Validation BCE Loss** | 0.0797 | 0.0936 |
| **Number of trainable parameters** | 94,331 | 182,075 |

**Figure 12.**
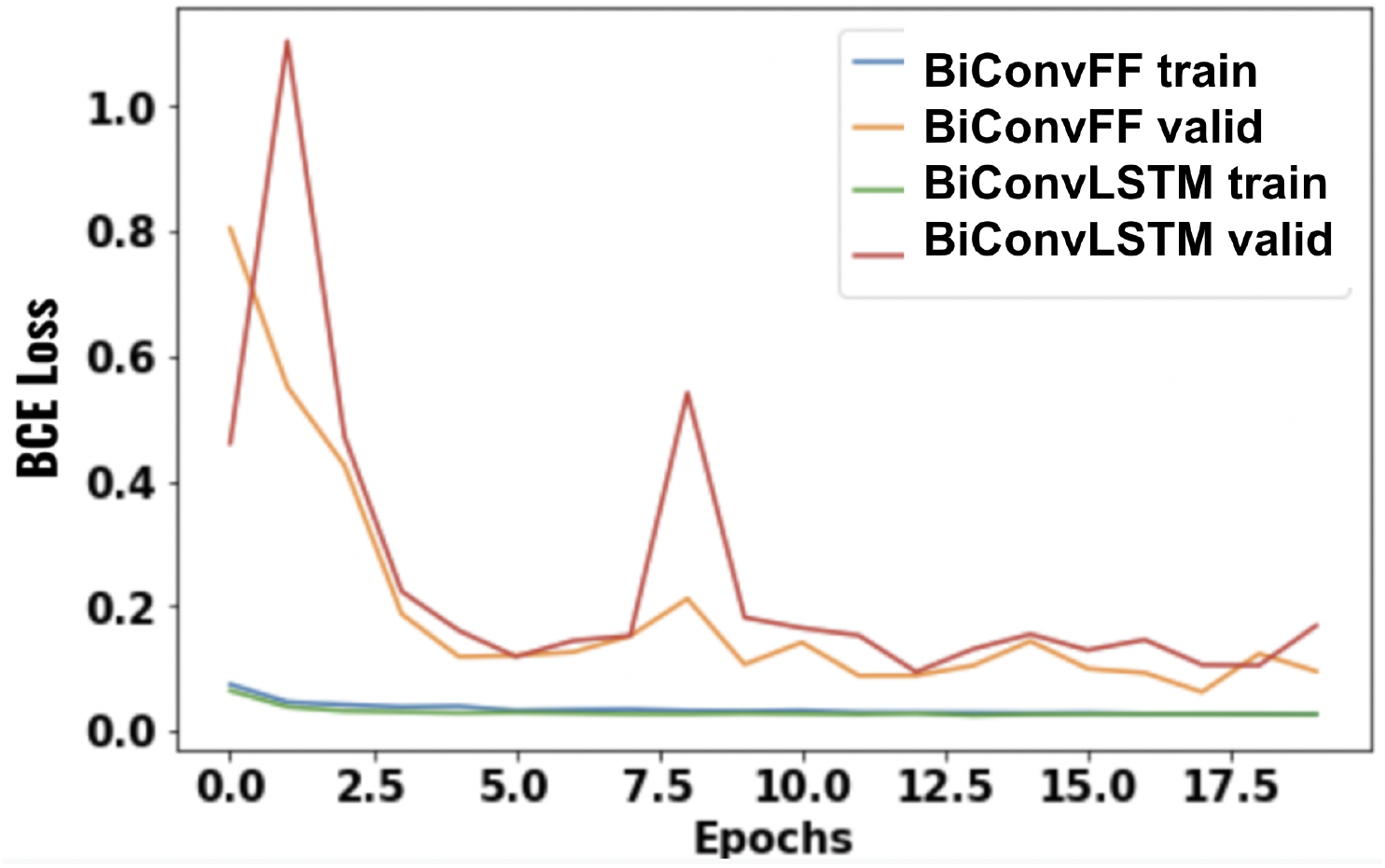
BiConvFFNN vs RNN-BiConvLSTM - Training and Validation Loss vs Epochs

**Figure 13.**
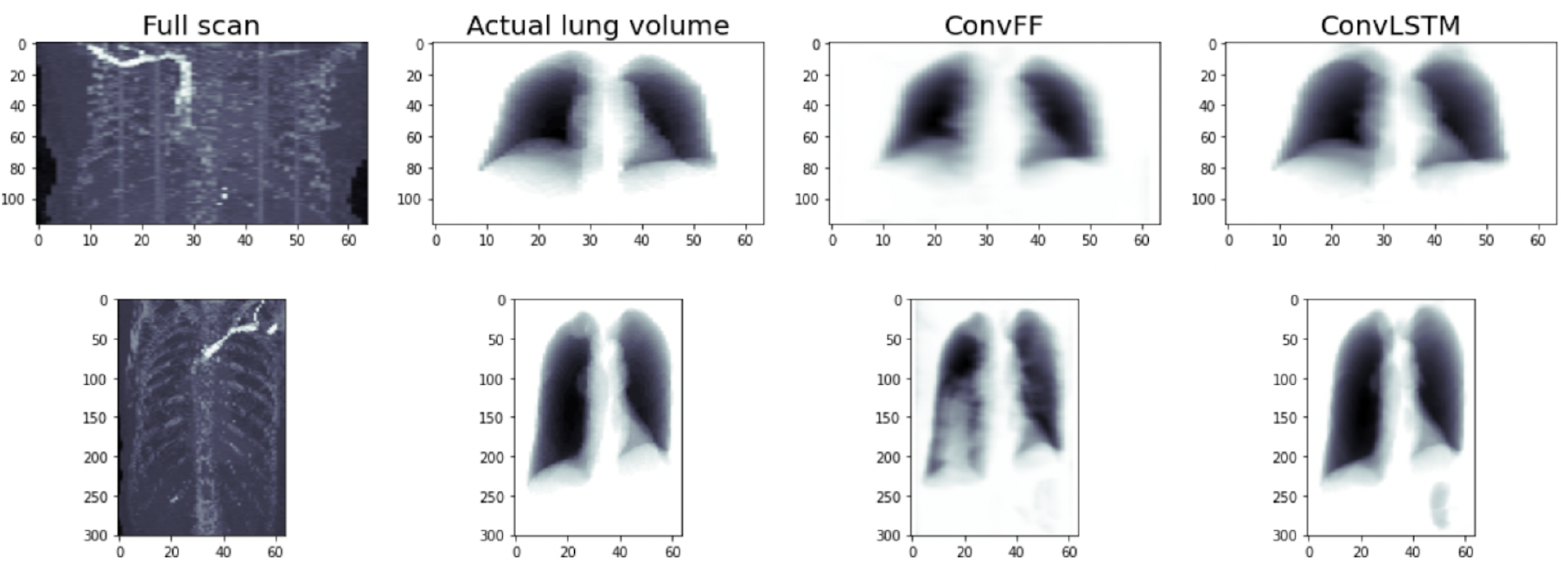
BiConvFFNN vs RNN-BiConvLSTM - Prediction results are shown in last two columns

#### Action recognition

Action recognition is the task of predicting the category of the action occurring in the video^20^. Each frame of the video contains the image contextual information of the action. The entire set of the frames in a video that depicts the action is fed as input to the neural network which predicts the category of the action. UCF11^54^ dataset is used for this task which contains 11 action categories: basketball shooting, biking/cycling, diving, golf swinging, horseback riding, soccer juggling, swinging, tennis swinging, trampoline jumping, volleyball spiking, and walking with a dog.

UCF 11 dataset is considered as one of the most challenging action recognition datasets due to its large variations in camera motion, object appearance and pose, object scale, viewpoint, cluttered background, illumination conditions, etc. In the dataset, a total of 1350 videos are split into 1200 videos in the training set, and 150 videos in the validation set. In view of the limitations in computational power, we reduced the frame size from 224 to 64 across both dimensions - width and height. The number of frames in each video is 30. To perform action recognition, a sequence of frames from the video is fed into the model.

The detailed layers of the proposed network are given in Table 11. The network is trained by backpropagating the cross-entropy loss between the softmax output class probabilities and ground truth class label using Adam optimizer for 10 epochs. We also trained the RNN-ConvLSTM network with the same training set. The number of neurons and the number of layers of the RNN-ConvLSTM network are the same as the ConvFFNN, except for the dropout. Dropout with a keep probability of 0.2 and 0.3 was applied after the ConvLSTM2D layer and the FC layer respectively in RNN-ConvLSTM. The RNN-ConvLSTM network is also trained for 10 epochs by using the same loss function and the same optimizer. The results from both networks are listed in Table 11. Trainable parameters are very close in this case because the FC layer connects 62× 62× 64 neurons (output from the last ConvFF2D/ConvLSTM2D layer) with 256 neurons and this alone takes 62, 980, 096 trainable parameters in both the networks. Therefore, number of trainable parameters taken by ConvFF2D layer is 77, 579 which is equal to 63, 060, 491 *−* (62, 980, 096 + 256 × 11) whereas number of parameters taken by ConvLSTM2D layer is 1, 54, 891 which is equal to 63, 137, 803 *−* (62, 980, 096 + 256 × 11). These numbers are shown in the Table 11. The accuracy of 99.4% and loss of 0.0015 achieved by ConvFFNN is significantly higher than the accuracy of 98.5% and loss of 0.0401 achieved by RNN-ConvLSTM network. Accuracy and loss with epochs by both the networks are shown in Figure 14, and 15 respectively.

**Table 11.** Network configuration of both ConvFFNN and RNN-ConvLSTM for action recognition and results in terms of accuracy, loss, and number of trainable parameters are shown here

|  | ConvFFNN | RNN-ConvLSTM |
| --- | --- | --- |
| **Network layers** | ConvFF2D, $3 \times 3 \times 64$<br>FC layer(256), ReLU<br>Softmax output layer(11) | ConvLSTM2D, $3 \times 3 \times 64$<br>Dropout(0.2)<br>FC layer(256), ReLU<br>Dropout(0.3)<br>Softmax output layer(11) |
| **Validation accuracy** | 99.4% | 98.5% |
| **Validation CE Loss** | 0.0015 | 0.0401 |
| **Number of trainable parameters** | 63,060,491 | 63,137,803 |

**Figure 14.**
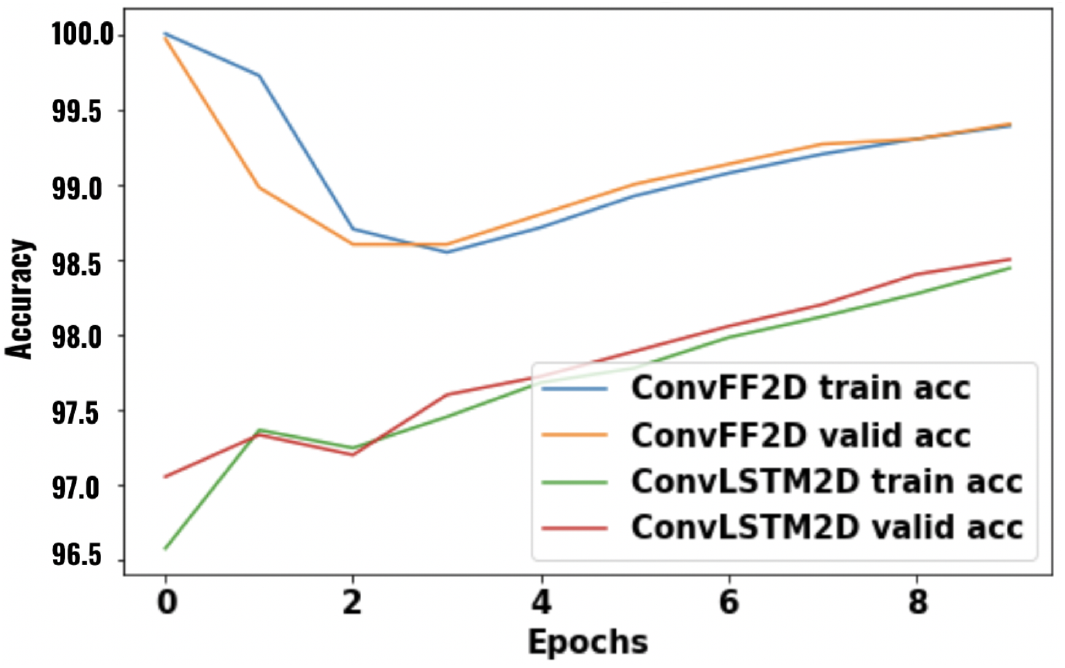
BiConvFFNN vs RNN-BiConvLSTM - Training and Validation Loss vs Epochs

**Figure 15.**
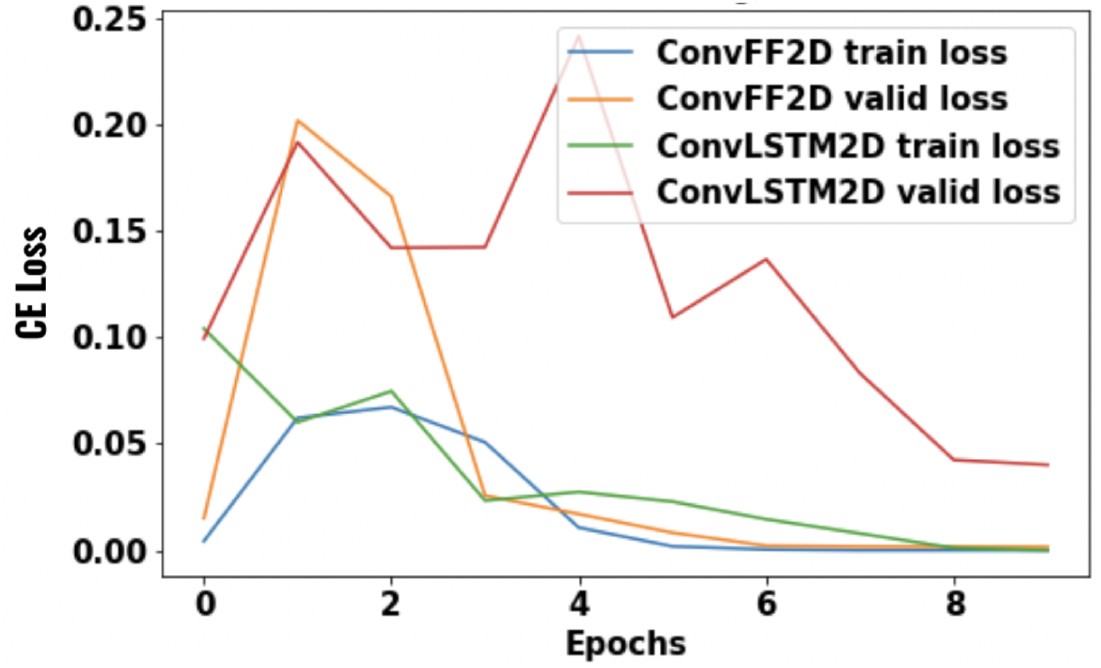
BiConvFFNN vs RNN-BiConvLSTM - Prediction results are shown in last two columns

### Hardware Benchmarking

To compare the performance of flip-flop network and RNN-LSTM in terms of hardware metrics in CPU, GPU and TPU, all the four models (FFNN, RNN-LSTM, ConvFFNN and RNN-ConvLSTM) were run for 1 epoch on a single data set. The details of the data and model architecture are listed in Table 12.

**Table 12.** Data and model architecture details used for hardware benchmarking

| Model | Data | Network Architecture |
| --- | --- | --- |
| **FFNN & RNN-LSTM** | Input Sequence length: 300<br>Input data dimension: 30<br>Output data dimension: 10<br>Batch size: 256 | FFNN/RNN-LSTM units: 10<br>Optimizer: SGD<br>Loss: MSE |
| **ConvFFNN & RNN-ConvLSTM** | Input Sequence length: 300<br>Input data dimension: $16 \times 16 \times 3$<br>Output data dimension: 30<br>Batch size: 64 | ConvFF/ConvLSTM: $3 \times 3 \times 8$, Flatten<br>Optimizer: SGD<br>Loss: MSE |
| **Validation accuracy** | 99.4% | 98.5% |
| **Validation CE Loss** | 0.0015 | 0.0401 |
| **Number of trainable parameters** | 63,060,491 | 63,137,803 |

The data is synthetically generated using Numpy’s random function to sample points from uniform distribution *U* (0, 1) and of data type float 64-bit. The benchmarking on CPU, GPU and TPU are carried out in Google Colaboratory free-tier. For CPU: Intel(R) Xeon(R) CPU @ 2.00GHz, GPU: Nvidia Tesla K80 (cuDNN version 8005 CUDAToolkit 11.2) and TPU: Colab TPU v2 are used for experimentation. Figure 16 and 17, show the result of memory requirements by the model to run one epoch. In GPU, the difference in memory requirements between flip-flop and RNN-LSTM based models are small because of Tensorflow automatically choosing cudNN based LSTM implementation to optimize but flip-flop runs on pure Tensorflow implementation^55^.

**Figure 16.**
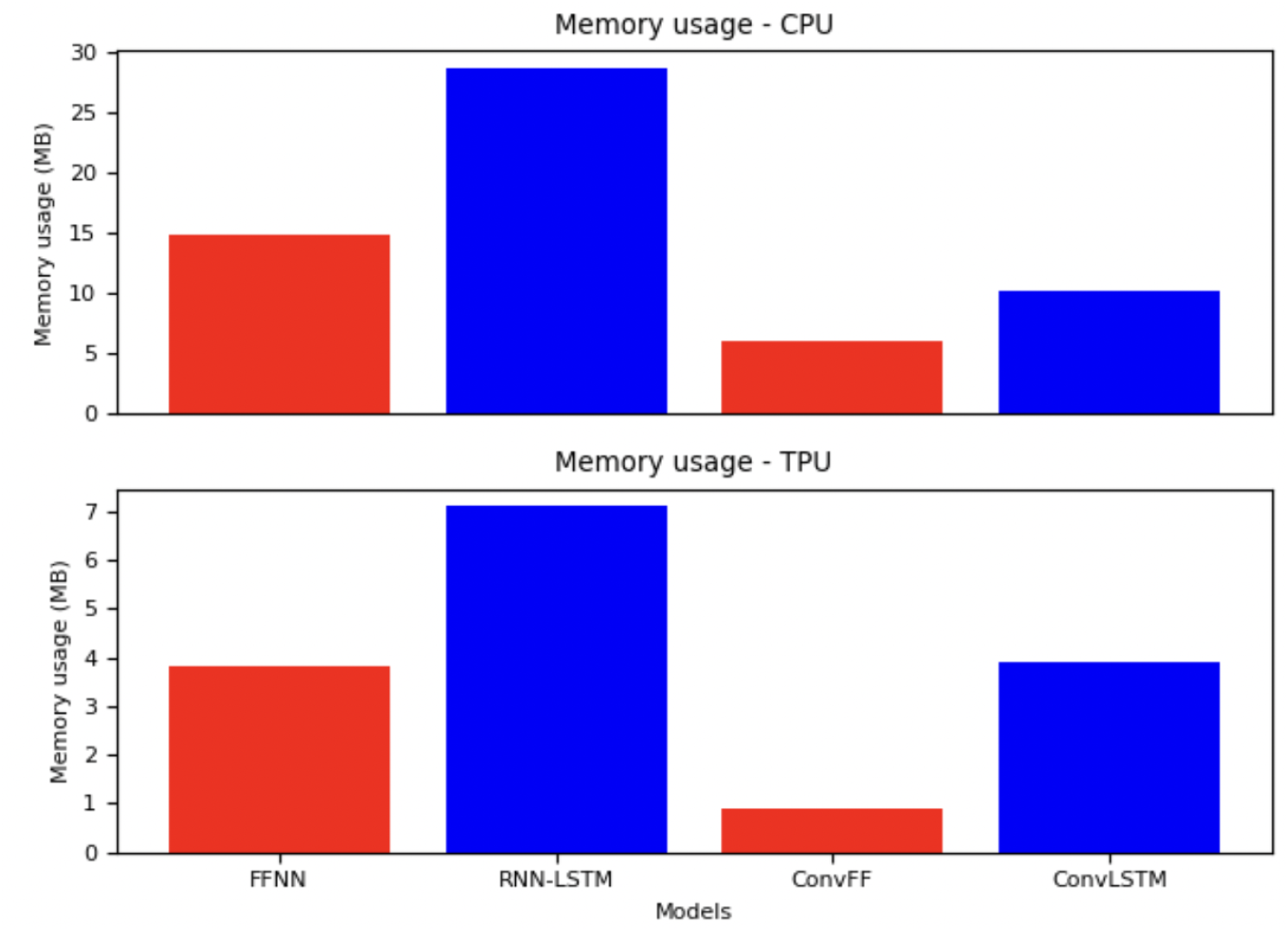
Memory usage of four models (FFNN, RNN-LSTM, ConvFF and ConvLSTM) in CPU, and TPU

**Figure 17.**
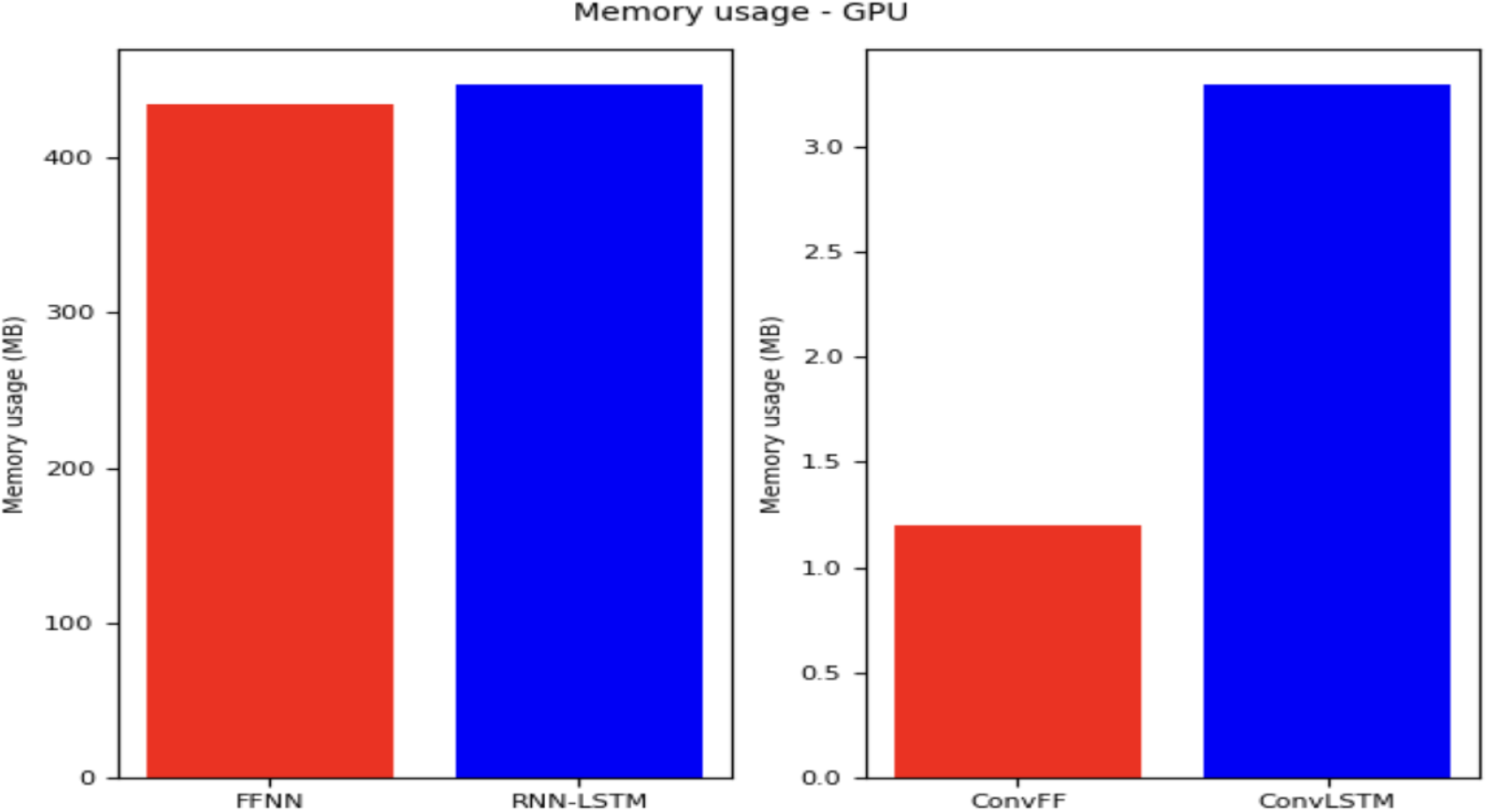
Memory usage of four models (FFNN, RNN-LSTM, ConvFF and ConvLSTM) in GPU

## Discussion

In the area of deep learning, in order to solve sequential decision making problems, several variations of RNNs have been proposed, out of which RNN-LSTM has been observed to perform the best most of the time. Four gates of LSTM that are required to enable it to remember/forget the (ir)relevant information in long-delay decision making problems, thereby making it computationally expensive. RNN-LSTM struggles to capture the long-term dependencies due to the problems of either gradient vanishing or gradient exploding. This gradient vanishing and exploding problems arise because of a large number of parameters that need to be updated in the LSTM cell.

The main reason for simulating the working memory in RNN is its resilience to noise and long time-lags between the stimulus and reward. Flip-flop networks solve this problem by exploiting the property of “store once retrieved forever”. Flip-flop neurons have inherent thresholding dynamics which make them resilient to noise within certain limits, a property called noise-margin in digital circuits. The proposed flip-flop neuron, which was inspired by the JK flip-flop, uses only two gates as opposed to four gates of LSTM, performs better or nearly the same as LSTM. We have analysed the results of all experiments and presented a Table 13 containing PLL, PAH, F:L, and the best performed neuron in each of the conducted experiments. Here, PLL is used to denote the percentage of loss (obtained by the best performed network either JK flip-flop deep network or LSTM deep network) that is lesser than the other network. PAH is used to denote the percentage of accuracy (obtained by the best performed netowrk) that is higher than the other network. F:L is the approximate ratio (flip-flop:LSTM) of required training parameters. Here, the F:L ratio in action recognition is considered the training parameters taken by flip-flop neuron and LSTM neuron alone (detail is given in action recognition section) respectively. In third row of the table, the word “Significantly” used to indicate that the network has outperformed with PLL more than or equal to 10 % and PAH more than or equal to 0.5 %.

**Table 13.** Summary table : Illustrates which of the neurons (JK flip-flop or LSTM) performed best in the seven experiments. D-1 and D-2 is Dataset-1 and Dataset-2 in Signal generation. PLL and PAH is percentage of loss and accuracy that is lesser and higher than the loss and accuracy of the beaten model. F:L is the ratio of required training parameters of flip-flop and LSTM neuron

| Task | Signal generation | Sentiment analysis | Handwriting generation | Text generation | Video frame prediction | Lung volume prediction in CT scan | Action recognition |
| --- | --- | --- | --- | --- | --- | --- | --- |
| **PLL** | 96.4 (D-1)<br>98.95 (D-2) | 4.53 | 29.62 | 54.22 | 78.58 | 14.8 | 96.25 |
| **PAH** | - | 0.014 | - | - | 4.34 | - | 0.9 |
| **F:L** | 1:2 | 1:2 | 1:2 | 1:2 | 1:2 | 1:2 | 1:2 |
| **Best performed by** | Significantly JK flip-flop | Insignificantly LSTM | Significantly JK flip-flop | Significantly JK flip-flop | Significantly JK flip-flop | Significantly JK flip-flop | Significantly JK flip-flop |

## Conclusion and Future work

In this study, it is shown that neural networks in which JK flip-flop neurons are incorporated are able to efficiently solve real-world sequential tasks of high difficulty level. Depending on what kind of data needs to be processed for a specific task, different versions of the flip-flop networks were modeled like FFNNs, BiFFNNs, ConvFFNNs, and BiConvFFNNs. We also derived learning rules for these networks by adapting from^21^. Comparison of both neurons on the seven experiments is summarized in Table 13. From the table we conclude that JK flip-flop neurons are more capable of remembering long sequences and perfectly able to manage the relevant and irrelevant information with a much smaller number of trainable parameters compared to LSTM.

There is still a need to explore more variations of the flip-flop neuron similar to the many variations of RNN-LSTM^44–51^ that are being developed to solve challenging sequential problems nowadays. Flip-flops can also be used to model cognitive tasks involving working memory (attributed to PFC’s up-down neurons) and related disease conditions^21^. The current flip-flop model can also be improved by taking inspiration from computational neuroscience to add additional sequence processing capabilities like chunking sequences^56^,^57^, working memory^58^,^59^, timing^60^ and processing hierarchical temporal representations^61^. The flip-flop neurons can also be combined with generative adversarial networks (GAN) for sequence generation tasks^50,51^. In the future work, we want to apply the flip-flop network to simulate human visual attention search and modelling very long sequences spanning more than 1000 time steps^62^.

## Acknowledgement

Sweta Kumari acknowledges the financial support of the Ministry of Human Resource Development for graduate assistantship. Vigneswaran Chandrasekaran acknowledges the financial support of the Indo-French Center for Promotion of Advanced Research (CEFIPRA) for graduate assistantship.

## Author contributions statement

The experiments were conducted by SK and VC both. SK wrote the main text. VSC contributed to providing the key ideas, editing the manuscript drafts, and providing insight into structure. All authors contributed to the article and approved the submitted version.

## Additional information

The authors declare that the research was conducted in the absence of any commercial or financial relationships that could be construed as a potential conflict of interest.

The corresponding author is responsible for submitting a competing interests statement on behalf of all authors of the paper.

### Quality Check

“This statement is to confirm that all methods were carried out in accordance with relevant guidelines and regulations “

## Notes

### Competing Interest Statement

The authors have declared no competing interest.

